# Nitrogen metabolic map of *Escherichia coli* mediated by multi-functional aminotransferases

**DOI:** 10.64898/2026.07.30.740019

**Authors:** Shogo Hataya, Mauricio Ferreira, Fayaz Soleymani, Sora Fukui, Marcos de V.V. Oliveira, Kaan Koper, Taichi E. Takasuka, Zoran Nikoloski, Hiroshi A. Maeda

**Author notes:** **Corresponding authors:** Taichi E. Takasuka, Zoran Nikoloski, Hiroshi A. Maeda.

## Abstract

Nitrogen is a key element of organic molecules essential to all life. Unlike extensively characterized carbon metabolic maps, it remains elusive how assimilated nitrogen flows through metabolic networks. Here we determined the nitrogen metabolic map of *Escherichia coli* and used it to construct an enzyme-constrained metabolic model that includes multi-functionality of aminotransferase enzymes responsible for nitrogen transfer reactions. We characterized substrate specificities of sixteen *E. coli* aminotransferase enzymes by evaluating 2,528 reactions, uncovering 56 previously unrecognized activities. The cellular concentrations of these aminotransferase enzymes were quantified and used to estimate their catalytic rates across all enzyme-substrate pairs, leading to improved predictions of nitrogen flows in *E. coli*. This work advances our fundamental understanding of the nitrogen metabolic map critical for metabolism and growth.

## Introduction

Nitrogen (N) is one of the essential elements of life as a key component for amino acids, proteins, nucleic acids, and other N-containing biomolecules. While assimilation and incorporation of N via the glutamine synthase-glutamate synthase (GS-GOGAT) and glutamate dehydrogenase (GDH) are thoroughly characterized (*1–4*), it remains poorly understood how assimilated N is distributed throughout the cellular metabolic network. This is in contrast to the extensively studied carbon (C) metabolic map of organisms (*5–8*). This lack in our fundamental understanding of N metabolism ultimately limits the ability to accurately predict N metabolic flows and design strategies to improve N use efficiency in any organism.

Aminotransferase (AT) family of enzymes transfer reduced N across different metabolic pathways and are central to the N metabolic network. These pyridoxal 5’-phosphate (PLP)-dependent enzymes evolved even before the origin of life (*9*, *10*) and catalyze transamination reactions between amino acid (amino group donor) and keto acid (amino group acceptor) substrates by a ping-pong bi-bi mechanism (*11*, *12*). A single AT enzyme can, in theory, mediate transamination reactions with over 380 substrate combinations, but their substrate specificities are still largely elusive, even in the model organism *Escherichia coli* (*10*, *13*). The functional annotations of the *E. coli* AT (EcAT) enzymes are based primarily on genetic analyses and limited biochemical data, which typically assess a specific substrate combination(s) of interest (*14–17*). Consequently, many other possible substrate combinations remain elusive for the majority of EcAT enzymes (*10*).

Prior works showed that many *EcAT* deletion strains do not necessarily show amino acid auxotrophy nor growth defects unless multiple *EcAT* genes are deleted (*18–20*). For example, L-aspartate (Asp) auxotrophy is achieved by knocking out *tyrB* and *aspC*, whereas an additional *ilvE* knockout is needed for L-phenylalanine (Phe), L-leucine (Leu), or L-isoleucine (Ile) auxotrophy (*18*, *21*). Consistent with these genetic studies, earlier biochemical analyses identified multi-substrate specificities of AspC, TyrB, and IlvE that utilize Asp and branched-chain amino acids (BCAAs), including Leu, Ile, and L-valine (Val) (*15*, *22–24*). In addition, we recently reported that many AT enzymes from a model plant, *Arabidopsis thaliana*, can utilize multiple substrates (*25*), which may also be the case in other organisms, including *E. coli*.

Defining the complete substrate specificities of AT family of enzymes is critical for determining a comprehensive structure of the N metabolic network, which can be achieved by employing genome-scale metabolic models (GEMs). These models entail a large-scale, *in silico* reconstruction of known metabolic reactions in an organism and allow simulation of metabolic phenotypes by calculating metabolic fluxes from reaction stoichiometry and the mass balance of metabolites (*26*). These simulations can be further improved by leveraging information on enzyme properties, e.g., the catalytic rate (*k_cat_*) of an enzyme for a particular substrate and enzyme concentration (*27*). These protein-constrained GEMs (pcGEMs) have been used to study diauxic growth (*28*), overflow metabolism (*29*), and have been a useful tool for predicting metabolic engineering strategies (*30*). However, catalytic rates are not available for many activities of multifunctional enzymes, like AT enzymes, preventing their use in the prediction of N flows through a modeled metabolic network.

To obtain N metabolic map of *E. coli*, here we first determined substrate specificity of sixteen EcAT enzymes encoded in the *E. coli* genome (*13*) using recently developed liquid chromatography-mass spectrometry (LC-MS)-based high-throughput AT substrate screening (*13*, *25*) followed by validation using on-target quantitative methods (**Fig. 1**). In addition, we quantified intracellular concentrations of sixteen EcAT enzymes using quantitative proteomics for the wild-type and four single *EcAT* gene deletion mutants. The enzymatic activity, intracellular enzyme concentration, and growth data were then integrated together into a pcGEM to estimate the *k_cat_* values of each EcAT; these were in turn used to refine a deep learning model for enzyme catalytic rates. This pipeline successfully improved the predictive capacity for AT substrate promiscuity. The results from the retrained *k_cat_* prediction model are then used to refine the predictions of N flow through the metabolic network of *E. coli* (**Fig. 1**).

**Fig. 1.**
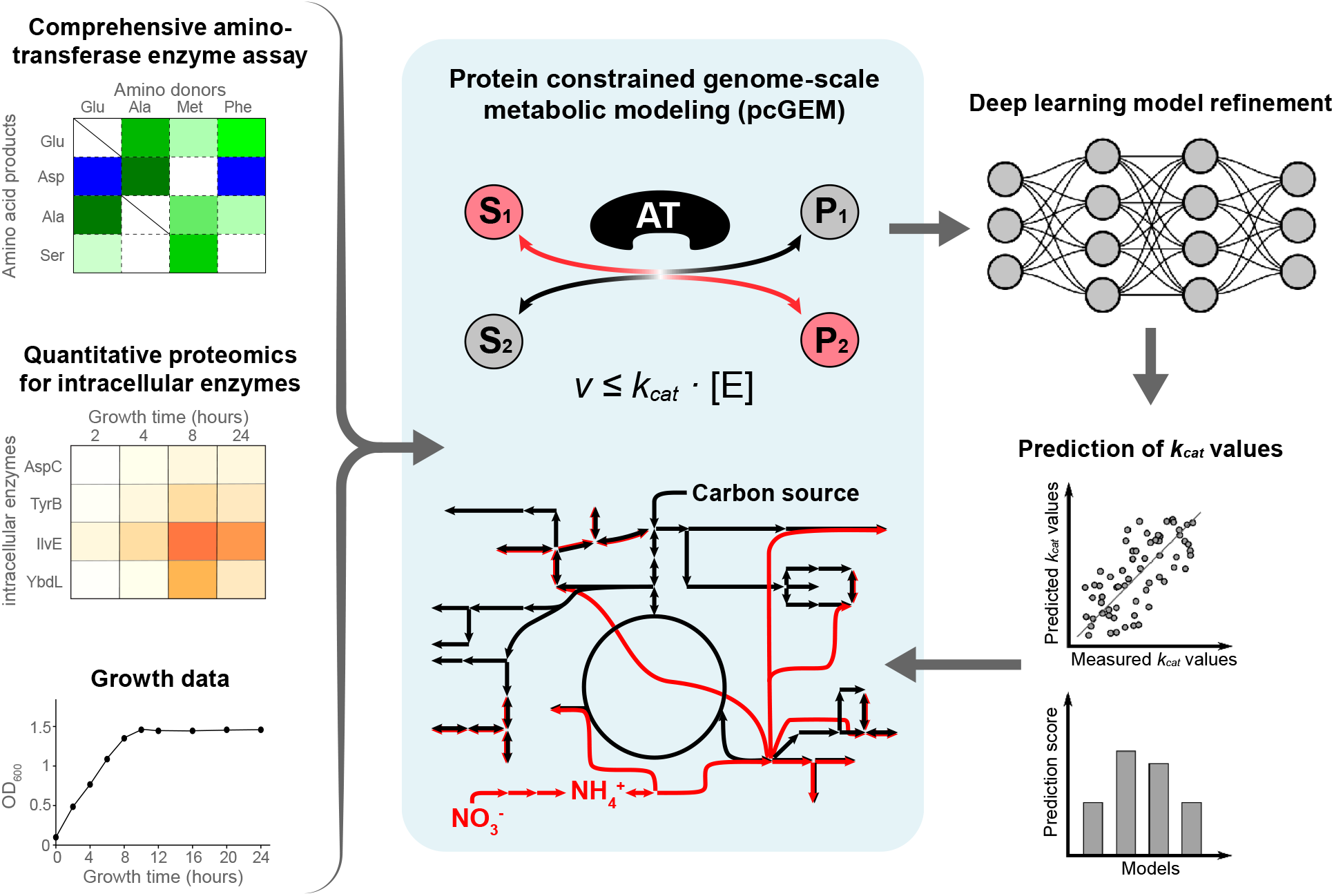
Construction of the *E. coli* nitrogen flux map. **(A)** Activities of *E. coli* aminotransferase (EcAT) enzymes, **(B)** their intracellular enzyme concentrations, and **(C)** growth measurements were determined by the EcAT substrate assays, quantitative proteomics, and growth assays, respectively. **(D)** The experimental data were integrated into a pcGEM of *E. coli*. Catalytic rates, i.e., *k_cat_* values, of EcAT enzymes were predicted using the proposed constraint-based modelling approach, kcatConv. **(E)** The predicted *k_cat_* were then integrated into the training data for the TurNuP *k_cat_* predicting model and retrained using the predicted *k_cat_* values to construct the *E. coli* nitrogen flux map.

## Results

### A comprehensive substrate mapping of sixteen AT enzymes of *E. coli*

To systematically screen substrate specificities of 16 EcAT enzymes across five AT classes (*10*), their corresponding genes were synthesized, recombinant enzymes were expressed and purified (**data S1**), and a LC-MS-based AT activity assay was conducted (**Fig. 2**, **data S2**) (*31*). Each enzyme was incubated with a reaction mixture containing relatively higher concentrations of substrates than the reported *K*_m_ of known ATs; one amino acid as the amino donor (5 mM) and a mixture of 15 biologically relevant keto acids (1 mM each) (*10*, *25*, *32*, *33*). The reactions were run to completion and % conversion of keto acids to amino acids were calculated. Each reaction was performed with one of the eight amino donors having different physico-chemical properties. Additionally, three different branched chain amino acids (BCAAs) were used as the amino donors to investigate additional enzymatic activities of TyrB, AvtA, and IlvE, which are known to be involved in the BCAA metabolism. In total, 2,528 unique reactions were conducted across the 16 EcATs, corresponding to 158 reactions per enzyme (**Fig. 2A, data S2**). The entire substrate mapping experiment was performed in two independent replicates, which yielded highly reproducible results (*r* = 0.954, *p-*value < 0.0001; **Fig. 2B**, **fig. S1**).

**Fig. 2.**
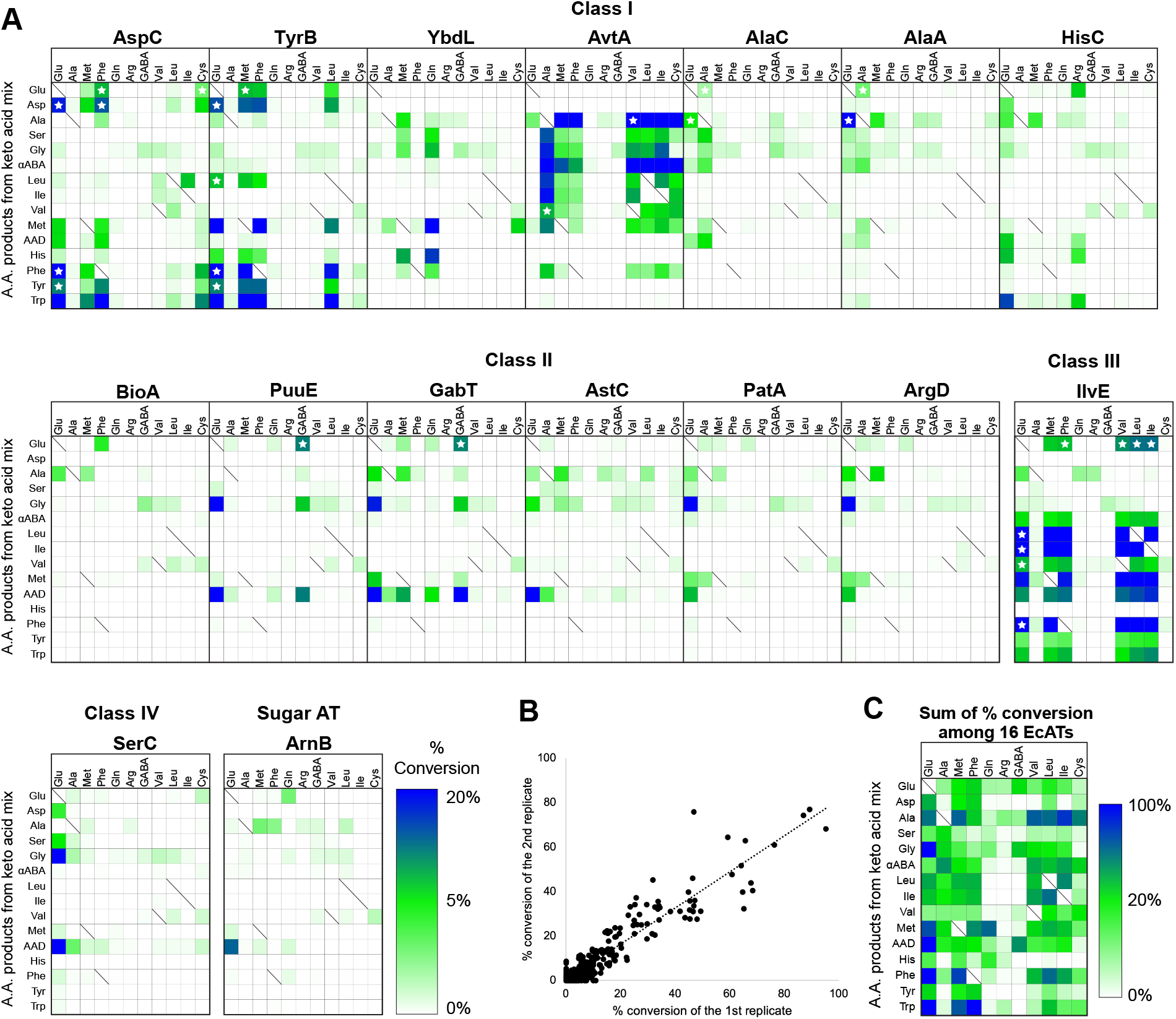
EcAT enzymes utilize combinations of multiple substrates. **(A)** Substrate specificity map of 16 EcATs across five phylogenetic classes, including Class I-IV and Sugar AT. The reactions were conducted using one amino donor at a time with a mixed 15 keto acids, and LC-MS was used to monitor the production of corresponding amino acid products. Reactions using the KRX strain carrying an empty vector were used as controls. The heatmap shows the % conversion by each enzyme from keto acid to amino acid, with the color bar indicating the degree of the % conversion. The % conversion values are the means of the two experimental replicates (**fig. S1**). The white stars denote previously reported activities. Abbreviations: αABA, α-aminobutyric acid; AAD, α-aminoadipate. **(B)** Correlation between the two independent replicates of the comprehensive EcAT substrate mapping experiment. Pearson’s correlation: *r* = 0.954, *p-*value < 0.0001. **(C)** The sum of the % conversion among the 16 EcATs.

Glu, a universal amino donor of many AT reactions (*10*, *34*), transaminated all 15 keto acids and could be produced by the reverse reaction with the corresponding substrate combinations by at least one EcAT (**Fig. 2C**). L-Alanine (Ala), L-methionine (Met), Phe, Val, Leu, Ile, and L-cysteine (Cys) also served as an amino donor to convert many keto acids in *E. coli*. All three BCAAs could be similarly used to synthesize most amino acids except for L-histidine (His). L-glutamine (Gln), L-arginine (Arg), and γ-aminobutyric acid (GABA) served as amino donors for synthesis of limited, specific amino acids (**Fig. 2C**).

To validate the LC-MS-based mapping results, we further conducted HPLC-based targeted and quantitative enzymatic assays on 59 selected reactions (**Table 1**), which include 44 previously unreported AT reactions that exceeded 20% conversion in the substrate mapping experiments (**Fig. 2A**). Each reaction was initiated by adding an individual EcAT enzyme, ranging from 0 to 10 mg/L, to a reaction mixture containing one specific keto acid and amino donor (see Methods). Overall, the HPLC assays validated 56 previously unreported activities (**Table 1**), as detailed in the next section, demonstrating that LC-MS-based substrate screening successfully captured comprehensive substrate specificity of 16 EcAT enzymes.

**Table 1.** Validated enzymatic activity of EcATs by HPLC-based assay. The % conversion is the average value of the 1^st^ and 2^nd^ replicates. Abbreviations: PPY, phenylpyruvate; IPyA, indole-3-pyruvic acid; 4MTOB, 4-methylthio-2-oxobutyrate; I5P, imidazole-5-yl-pyruvate; 4HPP, 4-hydroxyphenylpyruvate; 4MOP, 4-methyl-2-oxopentanoate; 3MOP, 3-methyl-2-oxopentanoate; AAD, α-aminoadipate; αABA, α-aminobutyric acid. N.D. denotes the products that could not be resolved on HPLC due to overlapping chromatographic peaks of an amino donor and an amino product (e.g., Val:4MTOB AT reaction catalyzed by IlvE). Specific activity represents the amount of amino acid (µM) produced per minute per enzyme concentration (mg/L). When the reaction became saturated within the tested conditions, their maximum detected activity is indicated. Stars indicate the reactions that were validated, whose activity plots are shown in **fig. S2**.

| Name | amino donor | keto acid | Amino acid product | % conversion | Note | Specific activity (μmol min<sup>-1</sup> mg protein<sup>-1</sup>) |
| --- | --- | --- | --- | --- | --- | --- |
| **AspC** | Glu | PPY | Phe | 36.6 | (12, 13) | >53.8* |
|  | Glu | IPyA | Trp | 64.3 | This study | >47.6* |
|  | Phe | IPyA | Trp | 55.8 | This study | 5.3* |
| **TyrB** | Glu | 4MTOB | Met | 31.5 | This study | 17.8* |
|  | Glu | IPyA | Trp | 48.7 | This study | 21.1* |
|  | Met | I5P | His | 4.3 | This study | 5.5* |
|  | Met | PPY | Phe | 38.5 | This study | 5.8* |
|  | Met | 4HPP | Tyr | 13.5 | This study | 2.7* |
|  | Met | IPyA | Trp | 52.2 | This study | N.D. |
|  | Phe | 4MTOB | Met | 26.1 | This study | >8.7* |
|  | Phe | IPyA | Trp | 54.4 | This study | >23.9* |
|  | Leu | PPY | Phe | 18.7 | This study | 1.2* |
|  | Leu | IPyA | Trp | 23.1 | This study | 1.3* |
| **YbdL** | Gln | 4MTOB | Met | 61.3 | This study | 5.8* |
| **AvtA** | Ala | glyoxylate | Gly | 18.0 | This study | 5.7 |
| | Ala | α-ketobutyrate | αABA | 42.3 | This study | >80.5 |
|  | Ala | 4MOP | Leu | 16.7 | This study | 2.9 |
|  | Ala | 3MOP | Ile | 19.1 | This study | 4.3 |
|  | Met | pyruvate | Ala | 31.9 | This study | 2.1 |
|  | Phe | pyruvate | Ala | 24.2 | This study | 1.2 |
| | Val | α-ketobutyrate | αABA | 35.9 | This study | 35.5 |
|  | Leu | pyruvate | Ala | 57.9 | This study | 6.8 |
| | Leu | α-ketobutyrate | αABA | 26.3 | This study | 7.1 |
|  | Ile | pyruvate | Ala | 81.7 | This study | 12.7 |
| | Ile | α-ketobutyrate | αABA | 37.8 | This study | 11.9 |
|  | Cys | pyruvate | Ala | 54.2 | This study | 4.6 |
| | Cys | α-ketobutyrate | αABA | 20.8 | This study | 4.9 |
| **PuuE** | Glu | glyoxylate | Gly | 20.8 | This study | 2.3* |
|  | Glu | oxoadipate | AAD | 46.2 | This study | 17.3* |
| | GABA | α-ketoglutarate | Glu | 12.0 | (16, 17) | >21.2* |
|  | GABA | oxoadipate | AAD | 12.1 | This study | 5.9* |
| **GabT** | Glu | glyoxylate | Gly | 18.3 | This study | 2.7* |
|  | Glu | oxoadipate | AAD | 80.7 | This study | 20.5* |
| | GABA | α-ketoglutarate | Glu | 11.6 | (18, 19) | >8.5* |
|  | GABA | oxoadipate | AAD | 38.5 | This study | 4.9* |
| **AstC** | Glu | oxoadipate | AAD | 83.1 | This study | 23.4* |
| **PatA** | Glu | glyoxylate | Gly | 29.7 | This study | 1.4* |
| **ArgD** | Glu | glyoxylate | Gly | 28.4 | This study | 1.5* |
| **IlvE** | Glu | 4MTOB | Met | 19.1 | This study | 7.6* |
|  | Met | 4MOP | Leu | 28.3 | This study | 6.8* |
|  | Met | 3MOP | Ile | 26.0 | This study | 5.8* |
|  | Met | PPY | Phe | 30.7 | This study | 6.1* |
| | Phe | α-ketobutyrate | αABA | 6.1 | This study | 2.0* |
|  | Phe | 4MOP | Leu | 32.8 | This study | 13.4* |
|  | Phe | 3MOP | Ile | 32.9 | This study | 10.6* |
|  | Phe | 4MTOB | Met | 18.7 | This study | 2.0* |
| | Val | α-ketobutyrate | αABA | 5.6 | This study | 2.2* |
|  | Val | 4MOP | Leu | 27.6 | This study | 37.4* |
|  | Val | 3MOP | Ile | 33.8 | This study | 29.3* |
|  | Val | 4MTOB | Met | 21.7 | This study | N.D. |
|  | Val | PPY | Phe | 34.6 | This study | N.D. |
|  | Leu | 3MOP | Ile | 61.9 | This study | N.D. |
|  | Leu | 4MTOB | Met | 24.7 | This study | 3.0* |
|  | Leu | PPY | Phe | 41.6 | This study | 21.2* |
|  | Ile | 4MOP | Leu | 36.5 | This study | N.D. |
|  | Ile | 4MTOB | Met | 26.3 | This study | 9.1* |
|  | Ile | PPY | Phe | 40.2 | This study | 20.7* |
| **SerC** | Glu | glyoxylate | Gly | 20.4 | This study | 2.2* |
|  | Glu | oxoadipate | AAD | 40.6 | This study | 15.3* |

### Multiple unreported activities were identified in *E. coli* AT enzymes

Among the Class I EcAT enzymes (*10*, *13*), AspC, TyrB, and AvtA showed broad substrate specificities compared to other Class I EcATs (**Fig. 2A**). Consistent with previous reports (*15*, *16*, *22*, *35–37*), marked with stars in **Fig. 2A**, our substrate mapping and HPLC-based assays (**Fig. 2**, **Table 1, fig. S2, data S2**) showed that AspC and TyrB can efficiently synthesize Asp, Phe, and L-tyrosine (Tyr) using Glu and aromatic amino acids (e.g., Phe) as the amino donor; in addition, AspC and TyrB exhibited weak activities to produce Glu using Met, Phe, Leu, or Cys amino donor. These assays also detected a number of previously unreported AT activities: AspC produced Trp using Glu, Met, Phe, or Cys amino donor and synthesized Met using Glu and Phe; similarly TyrB produced Met, Trp, His, Phe, and Tyr through Glu:4MTOB, Glu:indole-3-pyruvate (IPyA), Met:imidazole-5-yl-pyruvate (I5P), Met:PPY, Met:4-hydroxyphenylpyruvate (4HPP), Met:IPyA, Phe:4MTOB, Phe:IPyA, Leu:PPY, and Leu:IPyA AT activities (**Fig. 2**, **Table 1, fig. S2**). Overall, AspC and TyrB showed overlapping substrate specificities, consistent with their close phylogenetic relationships and three-dimensional structures (*10*, *38–40*), though TyrB can additionally use Met and Leu as an effective amino donor (**Fig. 2A**).

AvtA, known to interconvert Ala and Val (*20*, *22*, *41–43*), showed extremely broad substrate specificities: L-glycine (Gly), α-aminobutyric acid (αABA), Leu, and Ile could be produced efficiently (>15% conversion) when Ala was used as the amino donor, whereas Ala and αABA were generated using Met, Phe, Val, Leu, Ile, or Cys amino donor (**Fig. 2A**, **data S2**). HPLC-based assays further validated thirteen AvtA activities (**Table 1**), with Ala:α-ketobutyrate and Val:α-ketobutyrate as among the strongest AT activities (**Fig. 3A**). We also detected some reverse activities to those reactions—Ala:3-methyl-2-oxopentanoate (3MOP), Ala: 4-methyl-2-oxopentanoate (4MOP), Leu:pyruvate, and Ile:pyruvate AT activities (**Fig. 3A**)—to produce Ala and BCAAs. Therefore, these results revealed that, besides IlvE (*15*, *16*), AvtA contributes to the biosynthesis of all three BCAAs in *E. coli*, especially when Ala is available as an amino donor.

**Fig. 3.**
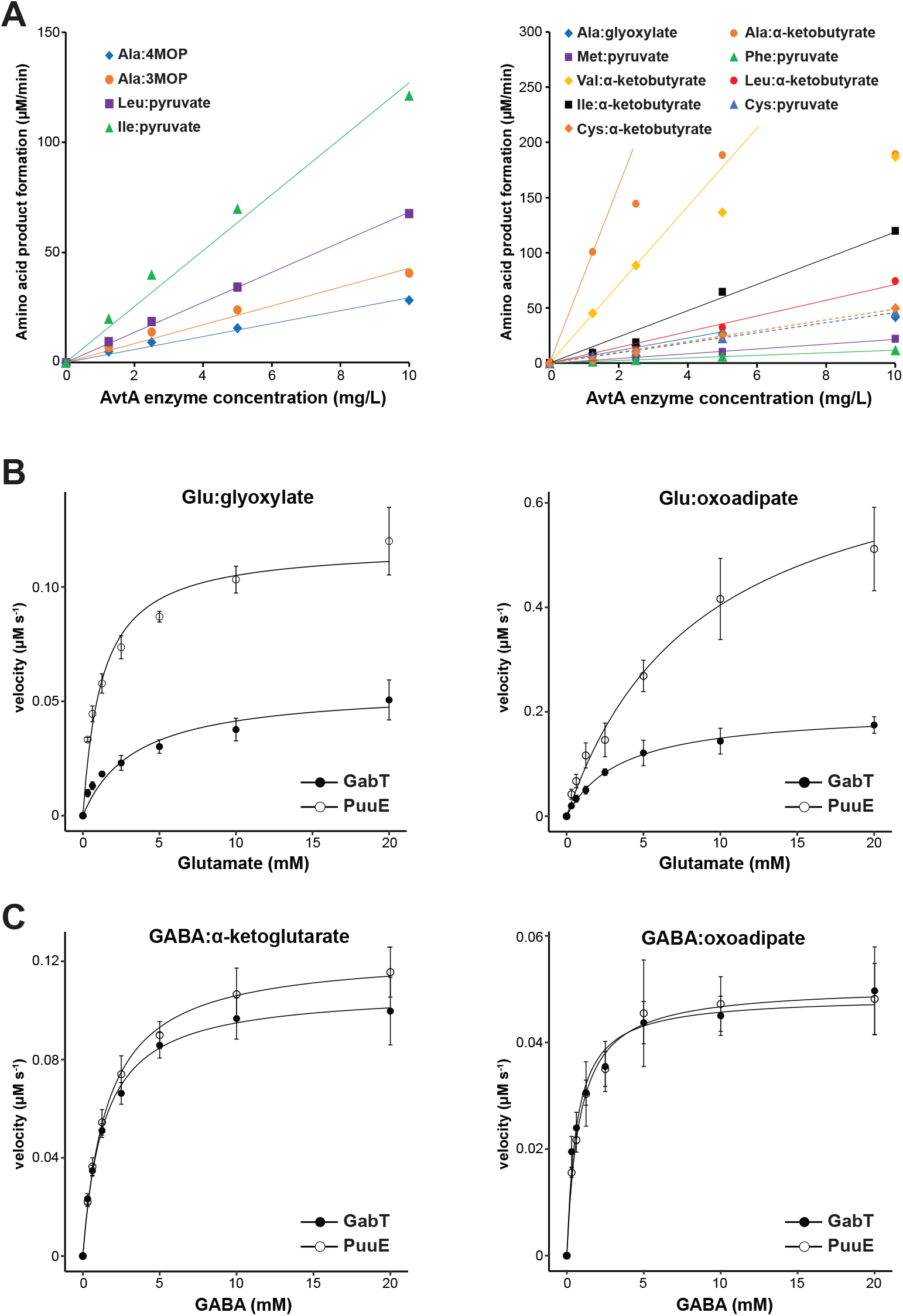
On-target quantitative AT activity assays validate multi-functionality of AvtA, GabT, an PuuE. **(A)** HPLC-based validation of AvtA activities for forward (left) and reverse (right) AT reactions between two branched chain amino acids: Leu and Ala (Leu:pyruvate and Ala:4MOP AT activities) and Ile and Ala (Ile:pyruvate and Ala:3MOP AT activities). Each reaction was performed once at four different enzyme concentrations. **(B)** Michaelis-Menten enzyme kinetic analysis of GabT and PuuE for Glu and glyoxylate (left) or oxoadipate (right). The AT activity of GabT and PuuE were tested with 1 mM keto acid and varying concentrations of Glu as substrates. GabT and PuuE plots are shown with green and red lines, respectively. Each data point is mean ± SD (n=3). **(C)** Kinetic analysis of GabT and PuuE for GABA and α-ketoglutarate (left) or oxoadipate (right). The AT activity of GabT and PuuE were tested with 1 mM keto acid and varying concentrations of GABA as substrates. GabT and PuuE plots are shown with green and red lines, respectively. Each plot is average of three replicates (n=3). Error bars show SD.

YbdL, a putative Met transaminase (*44*), catalyzed multiple AT reactions, which had not been previously recognized. YbdL showed efficient Gln:4MTOB and Gln:I5P AT activities to produce Met and His, suggesting that Gln is a suitable amino donor for YbdL in Met and His biosynthesis (**Fig. 2**, **Table 1, fig. S2**). YbdL also showed weaker AT reactions to produce Ala, L-serine (Ser), Gly, and Phe from the corresponding keto acids. These results suggest that YbdL has additional roles in the transamination of amino acids, especially in Met and His biosynthesis.

HisC is found within the His operon and exhibits Glu: L-histidinolphosphate AT activity (*45–49*). We found that HisC additionally has previously unreported activities to produce Trp, and to lesser extents His and α-aminoadipate (AAD), in the presence of Glu as the amino donor (**Fig. 2A**). The remaining Class I ATs, AlaC and AlaA, showed Glu:pyruvate AT activity (**Fig. 2A**, **data S2**) as previously reported (*20*), which is redundantly catalyzed by AvtA (*41*, *42*).

Among Class II ATs, PuuE and GabT showed GABA:α-ketoglutarate AT activity, as expected (*17*, *50–52*), but also exhibited strong Glu:oxoadipate activity to produce AAD (>45% conversion) and Glu:glyoxylate AT activity to produce Gly (∼20% conversion, **Fig. 2A, data S2, Table1, fig. S2**). Enzyme kinetic analyses (**Fig. 3B, C** and **Table 2**) further revealed that PuuE has comparable catalytic efficiency (*k*_cat_/*K*_m_) for Glu and GABA degradation (5.5-5.6 vs. 7.0-8.9 s^-1^mM^-1^, respectively) with glyoxylate, α-ketoglutarate or oxoadipate as the keto acid substrate. GabT showed somewhat higher catalytic efficiency for GABA (8.9-9.4 s^-1^ mM^-1^) than for Glu (1.2-3.4 s^-1^ mM^-1^; **Table 2**). Therefore, while GabT prefers GABA than Glu amino donor, PuuE can efficiently utilize both GABA and Glu for synthesizing AAD and Gly.

**Table 2.** Enzyme kinetic parameters of GabT and PuuE. The kinetic parameters, including *K_m_, V_max_, k_cat_,* and *k_cat_/K_m_*, of GabT and PuuE for Glu:glyoxylate, Glu:oxoadipate, GABA: α-ketoglutarate, and GABA:oxoadipate AT reactions. The kinetic parameters were determined by fitting the data to the Michaelis-Menten equation. Each value represents the mean of three independent replicates.

| enzyme | amino acid | keto acid (1 mM) | $K_m$ (mM) | $V_{max}$ ($\mu\text{mol s}^{-1}$) | $k_{cat}$ ($\text{s}^{-1}$) | $k_{cat}/K_m$ ($\text{s}^{-1} \text{mM}^{-1}$) |
| --- | --- | --- | --- | --- | --- | --- |
| **GabT** | Glu | glyoxylate | 3.43 | 0.056 | 3.25 | 1.20 |
|  | Glu | oxoadipate | 3.57 | 0.203 | 11.85 | 3.43 |
| | GABA | α-ketoglutarate | 1.37 | 0.108 | 12.62 | 9.43 |
|  | GABA | oxoadipate | 0.64 | 0.049 | 5.68 | 8.91 |
| **PuuE** | Glu | glyoxylate | 1.24 | 0.118 | 6.74 | 5.57 |
|  | Glu | oxoadipate | 8.69 | 0.756 | 43.16 | 5.49 |
| | GABA | α-ketoglutarate | 1.61 | 0.123 | 14.07 | 8.93 |
|  | GABA | oxoadipate | 0.82 | 0.051 | 5.77 | 7.00 |

BioA exhibited only weak activity for Phe:α-ketoglutarate, Glu:pyruvate, and Met:pyruvate AT activities (<5% conversion in the LC-MS mapping, **Fig. 2**, **data S2**), consistent with BioA’s reported main function as *S-*adenosyl-L-methionine (SAM): 8-amino-7-oxononanoate (AON) AT (*53*, *54*). Both AstC and ArgD were suggested to be involved in the ammonium-producing Arg succinyltransferase (AST) pathway and catalyze the *N2*-succinyl-L-ornithine:α-ketoglutarate and *N2*-acetyl-L-ornithine:α-ketoglutarate reactions, respectively, to produce Glu (*14*, *18*, *55–57*). However, the LC-MS mapping results showed that AstC and ArgD have Glu:oxoadipate and Glu:glyoxylate AT activities to produce AAD and Gly, respectively (**Fig. 2**, **data S2**), which were further verified by the HPLC-based assay (**Table1, fig. S2**). PatA is reported to be a putrescine:α-ketoglutarate AT responsible for putrescine catabolism (*58–60*). However, PatA also catalyzed Glu:glyoxylate AT reaction to produce Gly (**Fig. 2**, **Table 1, fig. S2**). Overall, among Class II EcATs, five enzymes (excluding BioA) exhibited AT activities to produce Gly and AAD as shared activities.

Among Class III AT enzymes, IlvE, a BCAA aminotransferase (BCAT) (*16*, *23*, *61*) showed previously reported activities of Glu:4MOP, Glu:3MOP, Glu:3-methyl-2-oxobutanoate (3MOB), Glu:phenylpyruvate (PPY), Met:α-ketoglutarate, Phe:α-ketoglutarate, Val:α-ketoglutarate, Leu:α-ketoglutarate, and Ile:α-ketoglutarate AT activities, to produce Leu, Ile, Val, Phe, and Glu, respectively (**Fig. 2**), as expected (*23*, *61*). We additionally detected 19 previously unreported activities: Glu:4MTOB, Met:4MOP, Met:3MOP, Met:PPY, Phe:α-ketobutyrate, Phe:4MOP, Phe:3MOP, Phe:4MTOB, Val:α-ketobutyrate, Val:4MOP, Val:3MOP, Val:4MTOB, Val:PPY, Leu:3MOP, Leu:4MTOB, Leu:PPY, Ile:4MOP, Ile:4MTOB, and Ile:PPY AT activities **Fig. 2**, **data S2**); 15 of them were further confirmed by the HPLC-based assays (**Table 1, fig. S2**). Therefore, IlvE not only functions as BCAT but can also exchange broad amino groups involved in Glu and Met, as well as in the reported Phe biosynthesis.

Among Class IV ATs, SerC is involved in pyridoxine and Ser biosynthesis by catalyzing the Glu:3-phosphohydroxypyruvate AT reaction to produce L-phosphoserine (*62–64*). However, we detected additional Glu:glyoxylate and Glu:oxoadipate AT activities at high efficiency (>20% conversion, **Fig. 2**, **data S2**), which were further verified by HPLC-based assay (**Table 1, fig. S2**). Therefore, SerC is potentially involved in the formation of Gly and AAD, besides Ser. The Sugar AT, ArnB, is known to function in the biosynthesis of a sugar nucleotide, particularly 4-amino-4-deoxy-L-arabinose (L-Ara4N) biosynthesis through the UDP-4-amino-4-deoxy-β-L-arabinose:α-ketoglutarate AT reaction (*65*, *66*). In addition, our analysis detected Glu:oxoadipate AT activity (∼14% conversion, **Fig. 2**, **data S2**).

Overall, our high-throughput LC-MS-based substrate mapping detected a large number of previously unreported AT activities (**Fig. 2**), many of which were further validated by on-target quantitative enzyme assays (**Table 1, fig. S2**). Therefore, these analyses together revealed that EcAT enzymes can catalyze many more AT reactions than previously reported.

### Quantification of EcATs intracellular protein levels to enable catalytic rate estimation

The identified substrate specificities of 16 EcAT enzymes enabled us to construct the N metabolic map of *E. coli*. Whilst capturing promiscuous activities of EcATs is rendered feasible in pcGEMs, this requires knowledge of catalytic activities that can be estimated using data on expression profiles and cell growth from different strains and/or environments (*27*). To this end, we quantified the concentrations of 16 EcAT enzymes at different growth stage by quantitative proteomics using isobaric tandem mass tag (TMT) (**Fig. 4A**). Cells were cultured separately in the LB medium and the M9 medium supplemented with glucose and BCAAs (hereafter simply M9 medium), which would support the growth of *ΔaspC* and *ΔilvE* mutants (*67*, *68*). The wild-type *E. coli* BW25113 strain started exponential growth at 2 hours and reached a plateau at 8 hours in the LB media, while in the M9 medium the cells started the log phase at around 12 hours and reached the stationary phase at around 24 hours (**Fig. 4B**). Thus, the proteomics analyses were conducted for the cells grown in LB medium for 2, 4, 8, and 24 hours (lag, log, end of log phase, and plateau phase, respectively), and cells grown in the M9 medium for 12 and 24 hours (the beginning and end of log phases, respectively). A duplex TMT-labeling proteomics was carried out for absolute quantification in *E. coli* cells, using cell-free-synthesized 16 EcATs with known concentrations as isobarically labeled protein standards with TMT^126^ (**fig. S3**). They were then mixed with the intracellular proteomes from the *E. coli* cells that were prepared at different time points and labeled with TMT^127^ (**Fig. 4A**).

**Fig. 4.**
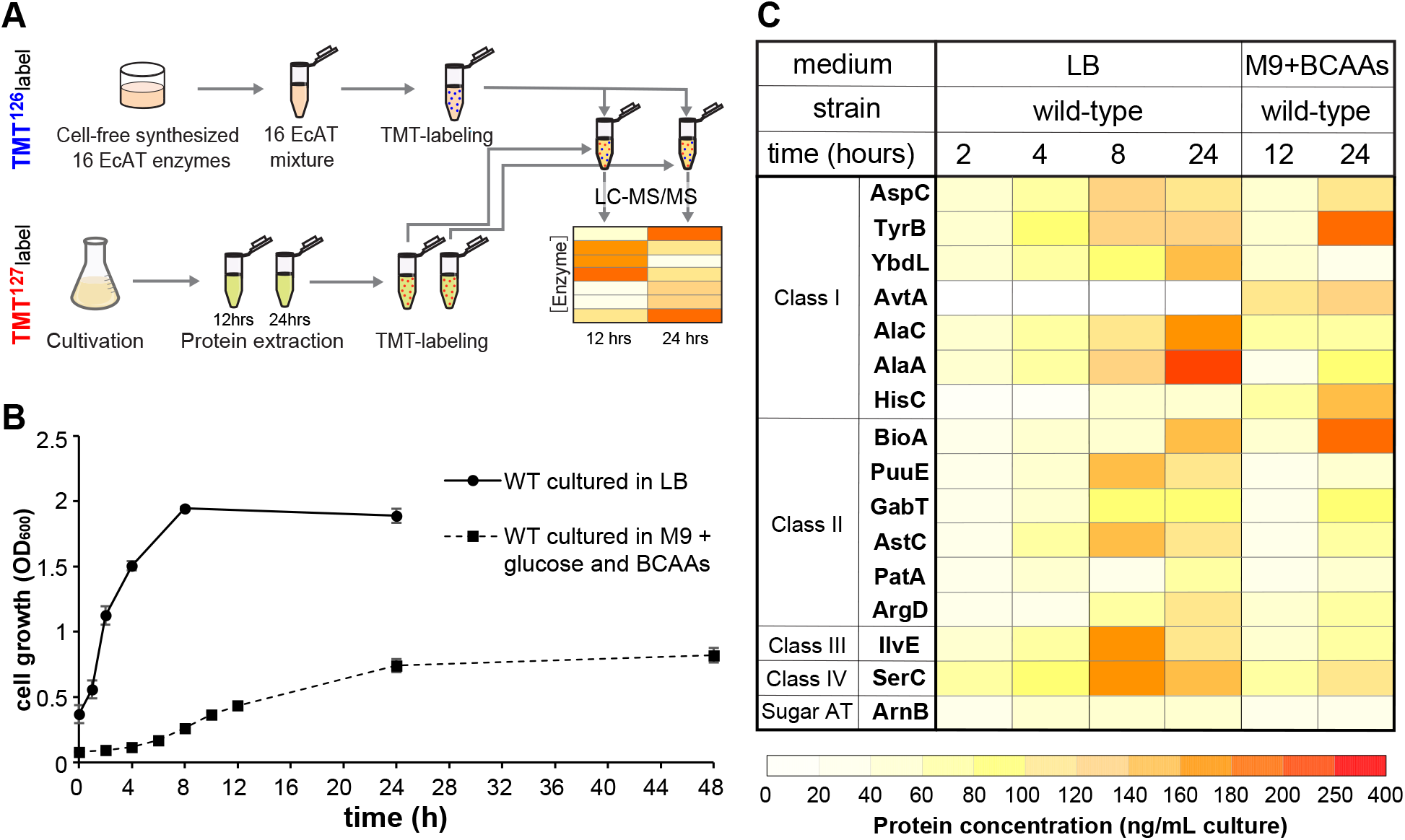
Intracellular changes in the proteome of EcAT enzymes. **(A)** Quantitative proteomic analysis for investigating time-dependent changes of 16 EcAT enzymes. **(B)** Growth curve of wild-type strain cultured in a rich LB medium or a minimal M9 medium supplemented with glucose and BCAAs (M9 medium). Each point is the average of separate growth and error bars show SD (n=3). **(C)** The protein concentrations of EcAT enzymes in the E. coli wild-type cells grown in LB and M9 media at each time course. The protein quantification was conducted by comparing the cell-free synthesized enzyme mixture from the representative dataset out of three biological replicates.

When the cells were grown in LB, all target enzymes except AvtA increased from the beginning (2 hours) to the middle (4 hours) of the log phase and reached the highest concentration at the end of the log (8 hours) or plateau phase (24 hours). The cells grown in the M9 medium took 12 and 24 hours to reach the intracellular enzyme levels similar to those of LB-grown cells at 4 and 8 hours, respectively (**Fig. 4B**). Most enzymes, except GabT, PatA, and ArnB, showed varied protein expression in the LB compared to the M9 media, whilst AvtA was detected only in the proteome datasets from the cells grown in the M9 medium (**Fig. 4C**). The expression patterns of AspC vs. TyrB (both in Class I), as well as PuuE vs. AstC (both in Class II) showed similar trends in both LB and M9 media. Although the concentrations of most of the enzymes were smaller than 100 ng/mL culture at the early log phase, they increased to or exceeded 100 ng/mL at the late log phase.

The duplex TMT labeling analyses were also conducted for four single knock-out mutants, *ΔaspC*, *ΔtyrB*, *ΔavtA*, and *ΔilvE*, to investigate how a deletion of these EcAT enzymes, showing a broad substrate specificity, affects other intracellular EcAT enzyme levels. When cultured in M9 medium without supplementation of BCAAs, Δ*aspC*, Δ*tyrB,* and *ΔavtA* displayed growth comparable to the wild-type, indicating that the absence of AspC, TyrB and AvtA is rescued by other EcATs (**fig. S4A**); in contrast, *ΔilvE* showed no growth. We found that *ΔtyrB*, *ΔavtA*, and *ΔilvE* strains showed similar levels of 16 EcAT expression, as compared to the wild-type (**fig. S4B**), consistent with TyrB, AvtA, and IlvE having functional redundancy in the BCAA biosynthesis (**Fig. 2**; **Table 1**). In contrast, *ΔaspC* showed reduced TyrB, BioA, PuuE, and IlvE, and increased AlaA, AstC, and PatA than wild-type cells (**fig. S4B**), suggesting that the function of AspC is partially insured by other EcAT enzymes. In the M9 media enriched with BCAAs, the lack of BCAA related enzyme did not affect the expression level of other EcATs except for YbdL, which was abundantly expressed in the *ΔaspC*, *ΔtyrB*, *ΔavtA*, and *ΔilvE* mutants, reflecting the broad substrate reactivities of YbdL (**Fig. 2**). Furthermore, these results suggested that the deficiency of one EcAT, and hence a specific amino acid, can alter the expression of other enzymes likely as a compensatory mechanism involving often multiple functionally redundant enzymes.

### Constraint-based modelling identifies catalytic rates of EcAT enzymes for multiple substrates

Towards constructing the N metabolic map of *E. coli*, the results of 2,528 reactions evaluated in this study (**Fig. 2**, **Table 1**) were added to the *E. coli* metabolic model iML1515 (*69*) to yield the eciML1515-EcAT model. This model was then constrained by integrating the absolute proteomics measurements of EcAT enzymes. The *k_cat_* values for the different substrate conversion reactions were considered as unknowns to be estimated using the relationship between fluxes, *k_cat_* values and protein abundance, whereby 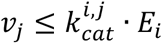 for the flux of reaction *j* catalyzed by enzyme *i* (see Material and Methods). We then integrated protein measurements at the 12-hour time point (end of log phase) into the model, in line with steady-state modelling, for five different strains, namely WT, Δ*tyrB*, Δ*avtA*, Δ*ilvE*, and Δ*aspC*, grown in the M9 medium. This constraint-based approach identified the minimum value for *k_cat_* capable of sustaining the reactions that are compatible with measured growth rates and protein abundance.

Since the constraint-based approach allows us to estimate *k_cat_* values, we first inspected the congruence of the estimated values for these parameters predicted by existing deep learning models. To this end, we compared the estimated *k_cat_* values obtained for each AT from DLKcat (*70*) and TurNuP (*71*). These deep learning tools require only structure-related features on substrates and products as well as sequence-related features on enzymes, and were trained on data on catalytic rates that are largely inferred from *in vitro* measurements. In contrast, the eciML1515-EcAT model fully leverages the physiological context and experimental data generated in this study to estimate *k_cat_* values. We found that the log-transformed pcGEM-estimated and DL-predicted catalytic rates for DLKcat and TurNuP yielded a small Pearson correlations (0.27 and 0.09, respectively; **fig. S5A, B**). Similar correlations were observed when comparing the predictions of *k_cat_* values from the DLKcat and TurNuP (**fig. S5C**). Next, the *k_cat_* values estimated using the eciML1515-EcAT model were used in refining the TurNuP model (**Fig. 5A)**. TurNuP was chosen over DLKcat given its better predictive capabilities, especially for non-canonical substrate-enzyme pairs (*71*). The Pearson and Spearman correlations between the log-transformed *k_cat_* values obtained using eciML1515-EcAT and the retrained TurNuP demonstrated significant improvement (Pearson correlation of 0.488 for 5-fold cross validated predictions) over the original model (**Fig. 5B**, **fig. S5**). Taken together, these results show that the metabolic context is important for prediction of enzyme kinetic parameters, with the integration of data-driven and physiology-informed modelling extending the capabilities of deep learning models.

**Fig. 5.**
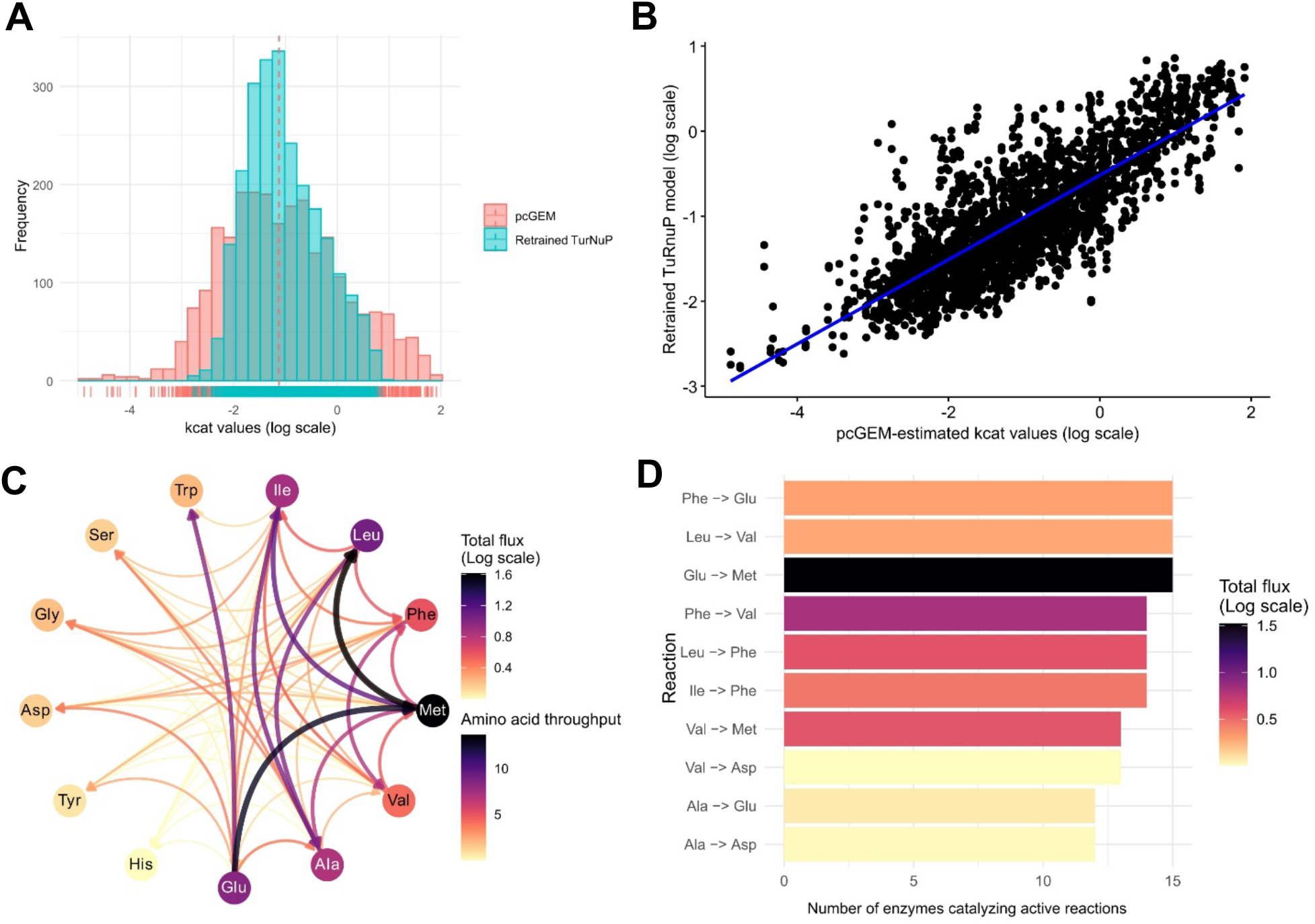
Performance of catalytic rate prediction and flux mapping of nitrogen metabolism. Following the estimation of *k_cat_* values using eciML1515-EcAT, the TurNuP model was retrained and *k_cat_* values were predicted. For eciML1515-EcAT, enzyme usage is encoded in the enzyme constraints and in the gene-protein-reaction (GPR) rules of each substrate conversion reaction. **(A)** Distribution of pcGEM-estimated *k_cat_* values and retrained TurNuP-predicted *k_cat_* values. y-axis denotes the frequency (i.e. proportion) of enzymes with a *k_cat_* value in a given bin. **(B)** Performance of the retrained TurNuP model. The retrained model shows improvement in predicting the *k_cat_* values of promiscuous enzymes, showing a Pearson correlation of 0.488, a mean squared error (MSE) of 1.0596, and a coefficient of determination (R²) of 0.232. Metrics were obtained for a 5-fold cross validation. **(C)** The total flux (represented in log scale) of individual edges is the sum of metabolic fluxes through all reactions for a specific substrate conversion catalyzed by different enzymes in the WT strain. Amino acid throughput for each node (i.e., amino acid substrate) represents the sum of the in-strength (inward edge weights) and the out-strength (outward edge weights) of all edges attached to that node. **(D)** For each substrate conversion reaction, multiple enzymes could be activated to catalyze the corresponding substrate conversion in *E. coli* WT (See the results of mutants in **fig. S6**).

### Constructing the *E. coli* N metabolic flux map

We next inspected the flux distribution across the eciML1515-EcAT model that is parameterized using the predicted *k_cat_* values from the refined TurNuP model. We simulated growth using flux balance analysis (FBA) and assessed how the flux is distributed across the substrate conversion reactions (**Fig. 5C, figs. S6A**, **S7A**). For comparison, when the conventional model iML1515 was also used to predict N fluxes, 2902 out of 2952 AT reactions were inactive (**fig. S8**). Therefore, our findings demonstrated that enzyme constraints allow us to study how promiscuous activities of EcAT enzymes contribute to the N flux of a given reaction in the N metabolic network, which could not be performed with the conventional constraint-based model.

The estimated N flux distribution, derived from the eciML1515-EcAT model, showed that Ala, Leu, Met and Phe, as hydrophobic amino acids, alongside with Glu form a core group of amino acids whose conversion reactions carry most of the N flux in the network, both in the WT (**Fig. 5C**) and the mutant strains (**fig S6, fig S7**). The most active conversion in the WT strain by total flux is Met to Leu (4.02 mmol/g_DW_/h), followed closely by Glu to Met (3.57 mmol/g_DW_/h). We also inspected the number of EcAT enzymes associated with reactions that carry flux and, thereby, contribute to the flux through each amino acid (**Fig. 5D**). We found that the flux through each amino acid in the core group identified above (Ala, Glu, Leu, Met and Phe) is supported by a low number of active AT enzymes catalyzing reactions of very high flux. For example, the Leu to Met conversion is mediated by three AT active enzymes, i.e., TyrB, AvtA, and IlvE, supporting flux-carrying reactions, while Met to Val is mediated by two such AT enzymes. The exceptions include the Glu to Met conversion (in the WT), mediated by 15 active enzymes (**Fig. 5D**).

Analyses of the Δ*tyrB*, Δ*avtA*, Δ*ilvE*, and Δ*aspC* mutants further showed substantial redistribution of N fluxes due to the loss of these individual enzymes. For example, Δ*tyrB* and Δ*ilvE* had a reduced flux from Glu to Met, consistent with the loss of *tyrB* and *ilvE*, both with strong Glu:4MTOB AT activity (**Fig. 2**). Instead, the most active conversions were shifted to Glu to Phe in the Δ*ilvE* (4.78 mmol/g_DW_/h) and Δ*avtA* strains (5.72 mmol/g_DW_/h) (**fig. S7D-E**), and to Phe to Val (8.81 mmol/g_DW_/h) in the Δ*tyrB* strain (**fig. S7C**). In line with the observed upregulation in the EcAT protein abundances in the Δ*aspC,* this strain exhibited the largest changes in N flux distribution among the four mutants, with the most active conversion of Ala to Met (9.28 mmol/g_DW_/h) (**fig. S6A-B**), resulting from the upregulation of AlaA and multiple other ATs (**fig. S3**). Like in WT, many AT reactions were still carried out by multiple enzymes. These findings suggest that the N metabolic network of *E. coli* WT functions to funnel amino groups primarily into the central Glu pool and redistributes them among various hydrophobic amino acids mediated by multiple AT enzymes. However, this N flux distribution pattern can flexibly change due to changes in abundance of certain AT enzyme having multi-substrate specificities.

## Discussion

The N metabolic map of *E. coli* constructed in this study reshapes our fundamentally understanding of the cellular N metabolic network. The functions of *E. coli* ATs have been extensively studied for many years through both genetic and biochemical analyses (*15*, *16*, *22*, *35*, *72*). However, our systematic re-characterization of EcAT enzymes, through LC-MS-based substrate mapping over 2,500 reactions of 16 EcAT enzymes (**Fig. 2**), followed by on-target quantitative enzyme assays (**Table 1**), detected a large number of previously unreported AT activities. For example, AvtA showed extremely broad substrate specificity to utilize Met, Phe, Val, Leu, Ile, and Cys in the presence of pyruvate, unraveling multi-functional capacities of EcAT enzymes. Conversely, multiple AT enzymes can redundantly catalyze a certain reaction, as demonstrated by both EcAT biochemical characterization (**Fig. 2**) and the metabolic modelling (**Fig. 5**). For instance, five Class II enzymes (i.e., PuuE, GabT, AstC, PatA, and ArgD) can redundantly catalyze the formation of Gly and AAD using Glu as the amino donor (**Fig. 2A**, **Table 1**). Although primary metabolic reactions are typically catalyzed by a highly specific enzyme with weak promiscuous activities if any (*73*, *74*), this study demonstrate that AT enzymes are multi-functional and can redundantly catalyze AT reactions across the cellular N metabolic network, depending on the cellular availability of different substrates.

The integration of proteomics measurements and growth data into the eciML1515-EcAT model further allowed estimation of catalytic rate; subsequent use of these rates improved predictions of metabolic phenotypes in comparison to the model that does not consider EcAT multi-functionality. Despite recent advances in predicting *k_cat_* values using deep learning tools (*75*), these models remain limited in their capabilities to accurately predict catalytic rates for enzyme-substrate pairs beyond the well-characterized pairs (*76*), available in databases such as BRENDA (*77*). In contrast, the combination of the pcGEM and deep learning models enabled us to refine the predictions of *k_cat_* values for the EcAT enzymes for different combinations of substrates and enzymes, beyond well-characterized enzyme-substrate pairs.

These AT multi-functionality and redundancy confer versatility and robustness to the N metabolic network, which are critical for organisms to ensure balanced synthesis of various N-containing compounds, including twenty proteogenic amino acids required for protein synthesis. Indeed, single *AT* gene deletion can be compensated by other ATs having overlapping activities (**Figs. 4**, **5**) (*16*, *20*, *61*, *68*). This versatility likely facilitated evolutionary diversification of AT enzymes, where orthologous ATs are not necessarily conserved across kingdoms, especially among microbes having extensive horizontal gene transfer events (*13*). Although N is essential for all organismal growth, both N input and output are dramatically different in different organisms. Some microbes can capture atmospheric dinitrogen (N_2_) to generate own ammonia (e.g., Rhizobia), while others must uptake N for their growth (e.g., *E. coli*, yeast, humans). Some organisms like plants often have limited N availability in the environments, even though they produce diverse N-containing natural products (e.g., alkaloids). Therefore, building up the current study, characterizing AT multi-substrate specificity and defining N metabolic map across diverse organisms will reveal how this essential metabolic network evolved and function across the Tree of Life.

## Materials and Methods

### EcAT enzyme expression and purification

The sixteen *EcAT* genes (**data S1**) were gene-synthesized at the Department of Energy (DOE), the Joint Genome Institute (JGI), and cloned into the pEU vector. EcAT enzyme-coding genes were amplified using primers listed in **data S3** and cloned into a modified pET28a vector (*31*) by sequence ligation-independent cloning (SLIC) or BsaI-based restriction enzyme digestion and ligation, accordingly to the manufacturer instructions (New England Biolabs, Inc., MA USA).

These 16 EcAT enzymes were recombinantly expressed and purified as previously described (*31*). Briefly, KRX *E. coli* cells (Promega, Madison, WI) harboring a modified pET plasmid with EcAT enzyme coding genes were grown on a Luria-Bertani (LB) agar (Sigma-Aldrich, St. Louis, MO) plate containing 50 µg/mL spectinomycin. Colonies were picked up and inoculated in 10 mL LB medium with 50 µg/mL spectinomycin, and incubated overnight at 37°C with shaking at 200 RPM. Subsequently, 10 mL of the culture was transferred to 1 L of Terrific Broth (TB) medium (Sigma-Aldrich) and grown at 37°C with shaking at 200 RPM until OD_600_ reaches ∼0.6. After setting the temperature at 22°C, 0.1% L-rhamnose and 0.15 mM isopropyl β-d-1-thiogalactopyranoside (IPTG) were added and the culture was incubated overnight at 22°C, 225 RPM. Cells were harvested by centrifugation at 6,000 *g* for 20 min at 4°C. The pellet was resuspended in 20 mL of lysis buffer containing 50 mM sodium phosphate (pH 8.0), 10% glycerol, 300 mM NaCl, 25 µM PLP (FUJIFILM Wako Pure Chemical Corporation, Osaka, Japan), and 0.25 mg/ml lysozyme (Sigma Aldrich). The cells were lysed by repeating freeze-thaw cycle three times, and the soluble part was collected by centrifugation at 18,000 *g* for 20-40 min at 4°C.

Cell pellet (from 40 mL culture) was resuspended in 2.4 mL BugBuster Master Mix (Merk Millipore) and lysed by incubation on a rotating mixer for 20 min. The cell lysate was centrifuged at 10,000 *g* for 30 min at 4°C and the supernatant was transferred to a new tube. Target proteins with His-tag were purified using nickel magnetic beads (Millipore), accordingly to a protocol provided by the manufacturer. Briefly, 100 µL of magnetic beads suspension was aliquoted to a 1.5 mL tube on a magnetic stand and magnetic beads were separated from the solution. The beads were equilibrated with 500 µL equilibration buffer containing 50 mM sodium phosphate (pH 8.0), 300 mM NaCl, 10% glycerol, and 25 µM PLP, twice. The equilibrated magnetic beads were resuspended in the soluble cell lysate, and the mixture was incubated for 30 min at 4°C with continuous mixing. Magnetic beads holding His-tagged proteins were separated from the solution on a magnetic rack and the supernatant was removed. The impurities were washed away using 500 µL wash buffer containing 50 mM sodium phosphate (pH 8.0), 300 mM NaCl, and 10 mM imidazole. The wash step was repeated three times. The washed beads were resuspended in 200 µL elusion buffer containing 50 mM sodium phosphate (pH 8.0), 300 mM NaCl, and 300 mM imidazole. After the incubation for 2 min, put the tube on a magnetic rack and collect the supernatant including recombinant His-tagged enzymes.

The purified proteins were desalted using 2 mL Zeba Spin Desalting Columns (Thermo Fisher Scientific) accordingly to manufacturer’s instruction. Briefly, storage solution in a desalting column set on a collection tube was removed by centrifugation at 1,000 *g* for 2 min. 1 mL desalting buffer containing 100 mM sodium phosphate (pH 8.0), 10% glycerol, and 25 µM PLP was poured to the column for equilibration. The buffer was removed by centrifugation at 1,000 *g* for 2 min. Equilibration was repeated three times. Equilibrated column was set on a new collection tube and purified proteins were applied on the center of the resin of the column. After the protein solution was fully absorbed, the proteins were collected by centrifugation at 1,000 *g* for 2 min. Protein concentration and purity was estimated by Bradford assay and SDS-PAGE analysis.

### AT substrate specificity mapping using LC-MS

To investigate substrate specificities of EcAT enzymes, LC-MS based high-throughput substrate mapping was performed. Enzyme reactions were performed for each purified EcAT in the reaction mixture containing 100 mM phosphate buffer pH 8.0, 1 mM ethylenediaminetetraacetic acid (EDTA), 25 µM PLP, 20 mg/L enzyme, 5 mM single amino acid, and 1 mM each of 15 keto acids: α-ketoglutarate, oxalacetate, pyruvate, β-hydroxypyruvate, glyoxylate, α-ketobutyrate, 4-methyl-2-oxopentanoate (4MOP), 3-methyl-2-oxopentanoate (3MOP), 3-methyl-2-oxobutanoate (3MOB), 4-methylthio-2-oxobutyrate (4MTOB), oxoadipate, imidazole-5-yl-pyruvate (I5P), phenylpyruvate (PPY), 4-hydroxyphenylpyruvate (4HPP), and indole-3-pyruvate (IPyA). To avoid substrate–product overlap, any keto acid that would generate the same amino acid used as the amino donor was excluded from the reaction mixture. For instance, α-ketoglutarate was omitted when Glu was used as the amino donor. Each reaction was initiated by mixing 10 µL of 200 mg/L single purified enzyme and 90 µL of the preheated reaction buffer, and the mixture was incubated at 37°C for 1 hour to reach the reaction equilibrium. The reaction was terminated by adding 400 µL LC-MS grade methanol and cooled on ice, followed by centrifugation at 15,000 *g* for 30 min at 4°C to precipitate insoluble chemicals. The supernatant containing amino acid products was collected.

Amino acids products were separated and detected by hydrophilic interaction chromatography using a HILIC-Z column (Agilent InfinityLab Poroshell 120 HILIC-Z, 2.1×150 mm, 2.7 µm) and Vanquish UHPLC system connected to Q Exactive Quadrupole-Orbitrap MS (Thermo Fisher Scientific). We run both negative and positive (only Gly) ionization mode using the settings as previously described (*31*). The mobile phases were: Buffer A (5 mM ammonium acetate and 0.2% acetic acid) and Buffer B (5 mM ammonium acetate, 0.2% acetic acid, and 95% acetonitrile). The sample was eluted at the flow rate of 0.45 mL/min with the following gradient of % buffer B: 100% from 0 to 1 min, 100% to 89% from 1 to 11 min, 89% to 70% from 11 to 15.75 min, 70% to 20% from 15.75 to 16.25 min, 20% from 16.25 to 18.5 min, 20% to 100% from 19.5 to 19.6 min, and 100% from 19.6 to 22.5 min. The mass spectrometry parameters for the Q Exactive system were configured in Full MS–ddMS² mode. The flow rates for sheath gas, auxiliary gas, and sweep gas were set to 55, 20, and 2, respectively, and the spray voltage, capillary temperature, and the S-lens RF level were set to 3 kV, 400°C, and 50, respectively. The Full MS settings included a resolution of 70,000, an AGC target of 3,000,000, a maximum IT of 100 milliseconds, a loop count of 2, a TopN of 2, an isolation window of 1 m/z, a stepped normalized collision energy (NCE) of 10, 20, and 40, and a spectrum data type set to centroid. For dd-MS² acquisition, the resolution was set to 17,500 with an AGC target of 100,000 and a maximum IT of 50 milliseconds. The loop count and TopN were both set to 2, with an isolation window of 1 m/z and stepped NCE values of 10, 20, and 40. The spectrum data type was also set to centroid. Standard curves for converting the peak area values to the concentrations were generated using authentic standards of amino acids. The % conversion values were calculated based on the molar ratio of the amino acid product to the initial keto acid. The extract of the KRX cells harboring an empty pET vector followed by the same purification as described above was used to subtract the background enzyme activity.

### EcAT enzyme activity validation using HPLC

For the HPLC-based enzyme assays, chemical reactions were carried out in 100 µL reaction mixture containing 100 mM sodium phosphate (pH 8.0), 1 mM EDTA (pH 8.0), 25 µM PLP, 5 mM single amino acid, 1 mM single keto acid, and 0, 1.25, 2.5, 5, or 10 mg/L EcAT enzyme. After incubating the solution at 37°C for 5 min, the reaction was terminated with 200 µL HPLC-grade methanol (Thermo Fisher Scientific). The samples were cooled at -20°C for 30 min and centrifuged at 21,000 *g* for 10 min at 4°C to precipitate insoluble compounds. Supernatant was transferred to a well of a 96-well plate and the amino acid product was analyzed by reverse phase HPLC using the ZORBAX eclipse-XDB C18 column (3 x 150 mm, 5 µm, Agilent) after derivatization by o-phthalaldehyde (OPA). The mobile phases were: Buffer A (0.05 M sodium acetate pH 5.88 and HPLC-grade methanol in a 95:5 ratio) and Buffer B (70% HPLC-grade methanol). Samples were eluted at the flow rate of 0.7 mL/min, the column temperature of 40°C, and a gradient elution of Buffer B: 0-1 min, 25%; 1-5 min, 25-80%; 5-10 min, 80%; 10-10.1 min, 80-100%; 10.1-12 min, 100%; 12-12.1 min 100-25%; 12.1-14 min, 25%. OPA-derivatized amino acids were detected by fluorescence detector (FLD, excitation 360 nm, emission 455 nm). The standard curves generated by varied concentrations of authentic amino acid standards were used to quantify the product formation, and specific enzyme activity (µmol min^-1^ mg protein^-1^) were calculated. In some cases, the HPLC-based analyses showed substantially higher activity (**Table 1, fig. S2**) than those detected by LC-MS-based screening (**Fig. 2**). For instance, TyrB catalyzed Met: I5P AT reaction more efficiently than Met:PPY and Met: 4HPP AT reactions by the HPLC-based assay (**Table 1, fig. S2**), whereas Met:I5P AT activity was much weaker than Met:PPY and Met:4HPP AT activity in the LC-MS screening. This is likely due to the presence of multiple substrates impacting certain reactions (e.g., competition) in the LC-MS screening, while the HPLC assays use specific substrate pairs.

### Enzyme kinetics analyses

The enzyme kinetic analyses of Glu:glyoxylate and Glu:oxoadipate AT activities were carried out in 100 µL reaction mixture containing 100 mM sodium phosphate (pH 8.0), 1 mM EDTA (pH 8.0), 25 µM PLP, 0, 0.3125, 0.625, 1.25, 2.5, 5, 10, or 20 mM of Glu, 1 mM single keto substrate glyoxylate or oxoadipate, and 2.5 mg/L GabT or PuuE. In the assay of substrate pairs GABA:α-ketoglutarate and GABA:oxoadipate, reaction mixture contained 100 mM sodium phosphate (pH 8.0), 1 mM EDTA (pH 8.0), 25 µM PLP, 0.3125, 0.625, 1.25, 2.5, 5, 10, or 20 mM of GABA, 1 mM single keto substrate α-ketoglutarate or oxoadipate, and 1.25 mg/L GabT or PuuE. The reactions were terminated by adding 200 µL HPLC-grade methanol after the reaction at 37°C for 5 min in a water bath. The samples were cooled at -20°C for 30 min and centrifuged at 21,000 *g* for 10 min at 4°C to precipitate insoluble compounds. Supernatant was transferred to a well of a 96-well plate and the amino acid product was analyzed by reverse phase HPLC using the ZORBAX eclipse-XDB C18 column (3 x 150 mm, 5 µm, Agilent) after derivatization by o-phthalaldehyde (OPA). The derivatized amino acids were detected by either of two methods. For the detection of Gly and AAD, the mobile phases were: Buffer A (0.05 M sodium acetate pH 5.88 and HPLC-grade methanol in a 95:5 ratio) and Buffer B (70% HPLC-grade methanol). Samples were eluted at the flow rate of 0.7 mL/min, the column temperature of 40°C, and a gradient elution of Buffer B: 0-1 min, 25%; 1-5 min, 25-80%; 5-10 min, 80%; 10-10.1 min, 80-100%; 10.1-12 min, 100%; 12-12.1 min 100-25%; 12.1-14 min, 25%. For the detection of Glu, the mobile phases were: Buffer A (0.1% ammonium acetate) and Buffer B (100% HPLC-grade methanol). Samples were eluted at the flow rate of 0.7 mL/min, the column temperature of 40°C, and a gradient elution of Buffer B: 0-3 min, 10-30%; 3-8 min, 30-50%; 8-9 min, 50-90%; 9-10 min, 90-10%; 10-12 min, 10%. OPA-derivatized amino acids were detected by FLD (excitation 360 nm, emission 455 nm). The enzymatic assay was conducted three times, and kinetic parameters were calculated based on the three replicates.

### Preparation of sixteen EcAT proteins for quantitative-protein mass spectrometry (qMS)

Cell-free translated EcAT enzyme mix (cell-free enzyme mix) was prepared by mixing equal amount of cell-free translated enzymes. All targeted enzymes synthesized by the wheat protein translation method were quantified by stain-free SDS-PAGE analysis (Bio-Rad Laboratories, California, USA). Expressed enzymes were detected and their concentrations were estimated by a Stain-Free imaging technology using TGX Stain-Free FastCast Acrylamide Kit (Bio-Rad Laboratories) and Gel Doc EZ Imager (Bio-Rad Laboratories) accordingly to the protocol provided by the manufacturer (Bio-Rad Laboratories). Briefly, 6 µL of translated samples were mixed with 2 µL of premixed 4x Laemmli protein sample buffer for SDS-PAGE (Bio-Rad Laboratories) and then heated at 95°C for 5 min. 6 µL of treated samples were run on the protein gel. The fluorescent signals detected by Stain-Free imaging technology rely on the number of Trp residues present in proteins on the gel. Thus, if there is no Trp residue in a targeted protein, the products must be stained by conventional gel staining methods, including Coomassie Brilliant Blue or fluorescent staining. The cell-free enzyme mix was made by mixing 306 µg of each enzyme and stored at -80°C until use. The cell-free enzyme mix contains a lot of wheat endogenous proteins, but these proteins do not affect proteomic analysis.

### Measurement of growth rate and cell dry weight of E. coli

After the preculture in 30 mL LB or M9 medium supplemented with 20 mM glucose and each 5 mM of BCAAs including Val, Leu, and Ile at 37°C, 160 RPM, overnight, absorbance was measured with Multiskan Go (Thermo Fisher Scientific, Illinois, USA). The preculture was added to a fresh LB or M9 medium to make 150 mL culture with OD_600_ at 0.1, and the cells grew at 37°C, 200 RPM for 48 hours. To investigate the growth rate, absorbance was measured at 600 nm every 2 hours until 12 hours after the start of incubation, and once more at 24 and 48 hours. For the cell dry weight measurement, cells were grown under the same condition and 5 mL of LB culture was collected at 2, 4, 8, and 24 hours, while 25 mL and 15 mL M9 culture was collected at 12 and 24 hours, respectively. Cells were precipitated by the centrifugation at 6,000 *g* for 10 min at 4°C. The supernatant was discarded, leaving 1 mL of liquid. The cell pellet was resuspended in the left liquid and transferred to a new 1.5 mL tube weighed in advance. After the cells dried at 30°C for several hours using an evaporator, weight was measured. To confirm that the cells are completely dry, weight was measured again next day.

### Cultivation and harvest of E. coli strains

Mutant strains were purchased from Keio-collection (**data S4**). Intracellular proteins at each growth level were extracted from *E. coli* cells and utilized for proteomic analysis after some treatments. BW25113 stain cells were collected from the glycerol stock and streaked on a LB agar plate followed by incubation overnight at 37°C. Single colonies were picked up from the plates and inoculated in 30 mL of LB liquid media. The *E. coli* cells were precultured overnight at 37°C with shaking at 160 RPM. 1 mL of the culture was put in a 3 mL glass cuvette, and the cell growth was checked by measuring the absorbance at 600 nm with Multiskan Go. The preculture was diluted with fresh LB liquid medium to adjust the OD_600_ to be 0.1, and the incubation at 37°C with shaking at 200 RPM was started. At each time point of 2, 4, 8, and 24 hours or 12 and 24 hours after the start of the incubation, 5, 15, 25 mL of the culture was collected to 50 mL FALCON tubes. The cells were harvested by the centrifugation at 6,000 *g* for 15 min at 4°C, and the cell pellet was stored at -80°C until use. Cell pellet was washed by resuspending in 100 µL of buffer A (20 mM phosphate buffer pH 7.4, 500 mM NaCl, 25 mM imidazole) and centrifuging for 3 min at 4,000 *g* at 4°C. Supernatant was discarded. The pellet was washed twice and then resuspended in 1 mL of buffer A. Cell suspension was sonicated, while being kept cool. Cell lysate was centrifuged at 16,000 *g* for 10 min at 4°C, and supernatant was collected in a new tube.

### Absolute quantification of cellular EcAT proteins by LC-MS/MS quantitative proteomics

Enzymes of interest extracted from *E. coli* cells were quantified by using isobaric Tandem Mass Tag (TMT) system (Thermo Fisher Scientific) (*78*). In this method, multiple samples can be detected simultaneously, and relative quantity of targeted proteins are estimated. By using a sample with known concentration as a standard, all detected targeted proteins can be absolutely quantified.

The amounts of protein in *E. coli* extract and the cell-free enzyme mix (16 ATs) were estimated by Bradford assay accordingly to the protocol provided by manufacturer (Bio-Rad Laboratories). Cell lysates and the EcAT mix were diluted with 100 mM TEAB (Thermo Fisher Scientific, Illinois, USA) to make 100 µL of 1 mg/mL protein solution. 5 µL of 200 mM TCEP (Thermo Fisher Scientific) was added to each solution, and samples were heated at 55°C for 1 hour to denature and cleave disulfide bonds. Sustain residues in samples were alkylated by adding 3.75 µL of 500 mM IAA (FUJIFILM Wako Pure Chemical Corporation) and incubated for 30 min in the dark. 100 µg of protein in each sample was precipitated by adding 600 µL of cold acetone (FUJIFILM Wako Pure Chemical Corporation) and incubated at -20°C overnight followed by the centrifugation at 8,000 *g* for 10 min at 4°C. The supernatant was discarded, and the pellet was air dried for 5-10 min. Precipitated protein was resuspended in 100 µL of 50 mM TEAB and digested into peptides by mixing with 20 µL of 0.125 mg/mL trypsin (Roche Diagnostics) and the incubation at 37°C overnight. Peptides were quantified using Pierce Quantitative Colorimetric Peptide Assay Kit (Thermo Fisher Scientific) accordingly the protocol provided by the manufacturer (Thermo Fisher Scientific) and diluted with 50 mM TEAB to adjust peptides concentration to be 1 mg/mL. Peptide quantification is required for proper TMT-labeling. Peptides were labeled using a TMTduplex Isobaric Label Reagents Set (Thermo Fisher Scientific) accordingly to the reported protocols (*79*) (Thermo Fisher Scientific). Briefly, TMT-labeling tags were dissolved in 41 µL of 100% acetonitrile (FUJIFILM Wako Pure Chemical Corporation) and mixed with sample peptides with 1:7 ratio of TMT-labeling reagent and sample solution followed by the incubation at RT for 1 hour. Cell-free translated enzyme mixture and *E. coli* extracted sample were labeled with TMT^2^-126 Label Reagent and TMT^2^-127 Label Reagent, respectively. Labeling reaction was stopped by adding 5% hydroxylamine (Thermo Fisher Scientific) and the incubation at RT for 15 min in the dark. 70 µL of TMT^2^-126 labeled cell-free synthesized enzyme mixture and 70 µL of TMT^2^-127 labeled *E. coli* extracted sample were combined to prepare a sample for quantitative MS analysis. Quantitative MS samples were evaporated at 30°C for 1 hour and resuspended in 20 µL of water followed by purification using ZipTip with 0.6 µL C_18_ resin (Millipore, Ireland). The solution X was prepared by mixing 95% of 0.1% formic acid in H_2_O and 5% of 0.1% formic acid in acetonitrile. The solution Y was also prepared by mixing 80% of 0.1% formic acid in acetonitrile and 20% of 0.1% formic acid in H_2_O. The ZipTip pipette column was equilibrated with 10 µL of the solution Y, and then 10 µL of X. 20 µL of the sample was pipetted approximately 20 times to attach peptides on the column. The column was washed 3 times with 10 µL of the solution X, and then peptides were eluted with10 µL the solution Y twice. The eluted sample was dried by a vacuum concentrator at 45°C for 20 min and resuspended in 12.0 µL of 0.1% formic acid in H_2_O. The sample peptides were separated and detected by using a nanoLC1000 (Thermo Fisher Scientific) hooked up with a hybrid linear ion trap-orbitrap mass spectrometer Q-Exactive Plus (Thermo Fisher Scientific). The MS results were analyzed by using Proteome Discoverer software v1.4 (Thermo Fisher Scientific). Operation parameters and data analysis conditions were set accordingly to an application note provided by the manufacturer (Thermo Fisher Scientific).

### Simulation of metabolic phenotypes and estimation of catalytic efficiencies of EcAT enzymes for multiple substrates

To assess how the different EcAT enzymes affect metabolism, especially nitrogen metabolism, the reactions corresponding to the 16 EcAT enzymes catalyzing the different substrate combinations were integrated into the *E. coli* metabolic model iML1515 (*69*), replicating the experimental design of the high-throughput substrate mapping. For each enzyme, one reaction was added for each combination of amino acid and keto acid substrates and resulting amino acid and keto acid products. Similar to the substrate mapping experiment, any keto acid that would generate the same amino acid used as the amino donor was excluded from the reaction mixture to avoid confounding results due to substrate–product overlap. Then, protein constraints were integrated into the model using the GECKO Toolbox 3 (*80*). The *k_cat_* values for all reactions except those of the included EcAT were predicted using DLKcat (*70*) and TurNuP (*71*). We have chosen these two DL models as DLKcat is the default *k_cat_* predictor integrated in the GECKO3 pcGEM reconstruction pipeline, and TurNuP covers many of the pitfalls DLKcat have (*81*) and holds up in performance compared to newer models (*82*). Each set of predicted *k_cat_* values was separately used to parameterize the curated iML1515 model, resulting in a DLKcat-parameterized protein-constrained model (pcGEM) and a TurNuP-parameterized pcGEM. Next, the pcGEMs were used to estimate *k_cat_* values of the enzymes for which no predictions from the deep learning (DL)-based models were employed in the optimization problem. The *k_cat_* values for these enzymes were integrated into the pcGEMs as unknowns, i.e., variables. The optimization problem was set up as follows:

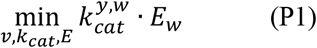

subject to:

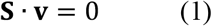

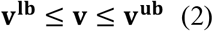

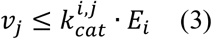

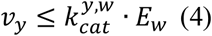

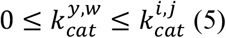

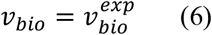

where *v_bio_* is the growth rate, **S** is the stoichiometric matrix, **v** is the flux distribution vector, **v^lb^** is the flux lower bound vector, **v^ub^** is the flux upper bound vector, *v*_j_ is the flux through reaction 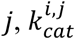 is the *k_cat_* value of enzyme *i* catalyzing reaction *j*, and *E_i_* is the concentration of enzyme *i*. (for the enzyme concentration, the absolute proteomics measurements of EcAT enzymes were used), *v_y_* is the flux through the substrate conversion reaction 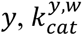 is the *k_cat_* value of enzyme *w* catalyzing the substrate conversion reaction 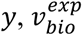 is the growth rate experimentally determined from the growth curves. The set of enzymes *w* is a subset of *j*, referring to the enzymes that also catalyze the added substrate conversion reactions *y* in addition to their native reactions *i* in the metabolic network. Given that the substrate affinity is lower for the non-native enzyme-substrate pairs evaluated in the simulation, the *k_cat_* values for these enzymes in the substrate conversion reactions are assumed to be lower than the *k_cat_* values for these same enzymes when catalyzing their native reactions, captured in Eq. (5).

To ensure that the estimated *k_cat_* values are consistent across the strains used for the experiments, the absolute proteomics measurements of the EcAT enzymes in different strains at the 12h time point were integrated into the pcGEM. Then, each resulting proteomics-constrained pcGEM was concatenated into a single model, such that the optimization simultaneously considered both wild-type (WT) and KO strains in the same linear problem. To achieve this, the stoichiometric matrix and all vectors (right-hand side, fluxes, reaction bounds, objectives) of each individual pcGEM were concatenated to form one single stoichiometric matrix and one of each vector, which contains the distinct mass balance and stoichiometric constraints of all strains. The introduced *k_cat_* variables for the substrate conversion reactions were kept the same for all strains, ensuring consistency across experimental conditions.

All code was written in MATLAB (The MathWorks Inc., Natick, Massachusetts), using functions from the packages COBRA Toolbox 3 (*83*), RAVEN Toolbox 2 (*84*), and GECKO Toolbox 3 (*80*). All optimization problems were set up and solved using Gurobi (*85*).

The *k_cat_* values estimated by the constrained-based approach were used to refine the *k_cat_* prediction tool TurNuP. They were integrated into its training set, and the deep learning model was retrained following the methods described in (*71*). The retrained model was then used to predict *k_cat_* values for the enzymes included in the pcGEM version of iML1515.

Lastly, to simulate growth and assess the flux distribution across the metabolic network, the predicted *k_cat_* values from the refined TurNuP model were integrated into the pcGEM. Then, flux balance analysis (FBA) with protein constraints was performed:

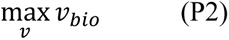

subject to:

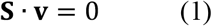

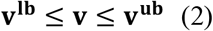

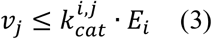

For the conventional iML1515 model, we performed the same optimization, albeit without the enzyme constraints as represented in Eq. (3).

## Acknowledgements

This work was supported by the US Department of Energy (DOE), Office of Science, Office of Biological and Environmental Research, Genomic Science Program (DE-SC0020390) to T.E.T., Z.N., and H.A.M., the Joint Genome Institute (JGI) award no. CSP-503757 to T.E.T. and H.A.M., the U.S. NSF PGRP award IOS-2312181 to H.A.M as well as the ALFAFUELS project funded by the European Union under Grant Agreement Number 191122224 and the German Research Foundation (DFG) project no. NI 1472/16-1 to Z.N, and the JSPS KAKENHI Grant Number 23K27036, 25K22375, and JST ALCA-Next (FS) Grant Number JPMJAN24D6 to T.E.T. S.H. was supported by Hokkaido University–Hitachi Joint Cooperative Support Program for Education and Research. The AT gene synthesis was carried out as a part of the CSP-503757 project by Drs. Yasuo Yoshikuni, Jan-Fang Cheng, Miranda Harmon-Smith, Sangeeta Nath, and Angela Tarver at the DOE JGI, a DOE Office of Science User Facility, which is supported by the DOE Office of Science operated under Contract No. DE-AC02-05CH11231.

## Author contributions

Conceptualization: TET, ZN, HAM

Methodology: MO, KK, MF, ZN

Investigation: SH, MF, FS, SF, MO

Visualization: SH, MF, FS, SF, HAM

Funding acquisition: TET, ZN, HAM

Project administration: TET, ZN, HAM

Supervision: TET, ZN, HAM

Writing – original draft: SH, MF, TET, ZN, HAM

Writing – review & editing: SH, MF, FS, SF, MO, KK, TET, ZN, HAM

## Competing interests

Authors declare that they have no competing interests.

## Data, code and materials availability

All data are available in the main text or the supplementary materials. All code used in this study is available at: https://github.com/mauricioamf/Ecoli_AT_analysis. The expression plasmids for expressing all EcATs are available upon request through a simple materials transfer agreement (MTA).

## Notes

### Competing Interest Statement

The authors have declared no competing interest.

