## Supplementary_Materials for "Nitrogen metabolic map of *Escherichia coli* mediated by multi-functional aminotransferases"

**Supplementary Figures:**

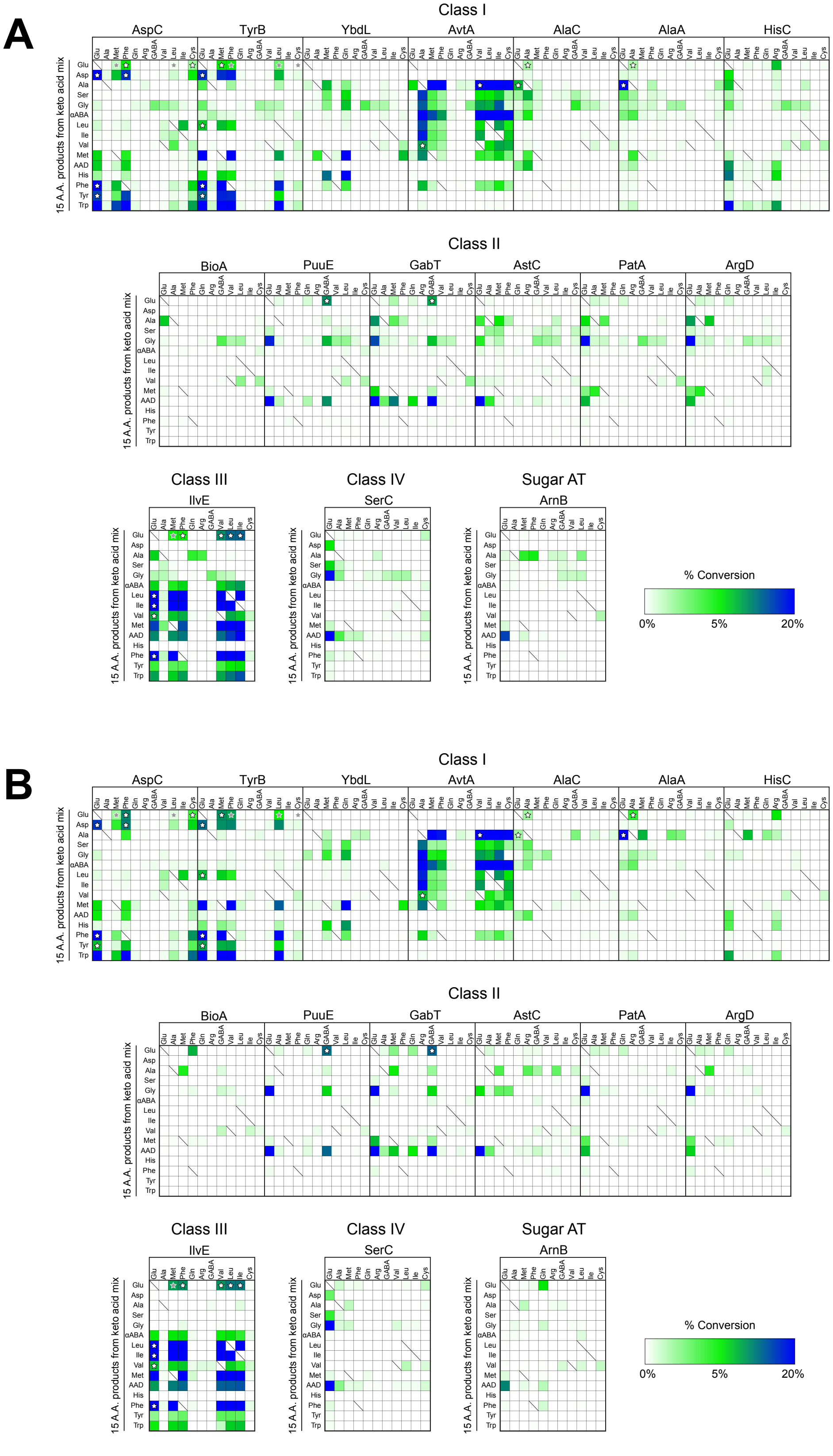

**Supplementary fig. S1. High reproducibility between the first and second runs of EcAT substrate screening experiments**

**(A)** The first and **(B)** the second replicates of the substrate mapping of 16 EcATs across five phylogenetic classes, respectively. The reactions were conducted using one amino donor and 15 keto acids as substrates, and the amino acid products were detected by LC-MS/MS. The heatmap shows the % conversion from keto acid to amino acid, as indicated in the color scales. As in **Fig. 2A**, the white and gray stars denote previously reported and indirectly suggested activities, respectively. The average % conversion values of the two replicates were calculated to generate **Fig. 2A**.

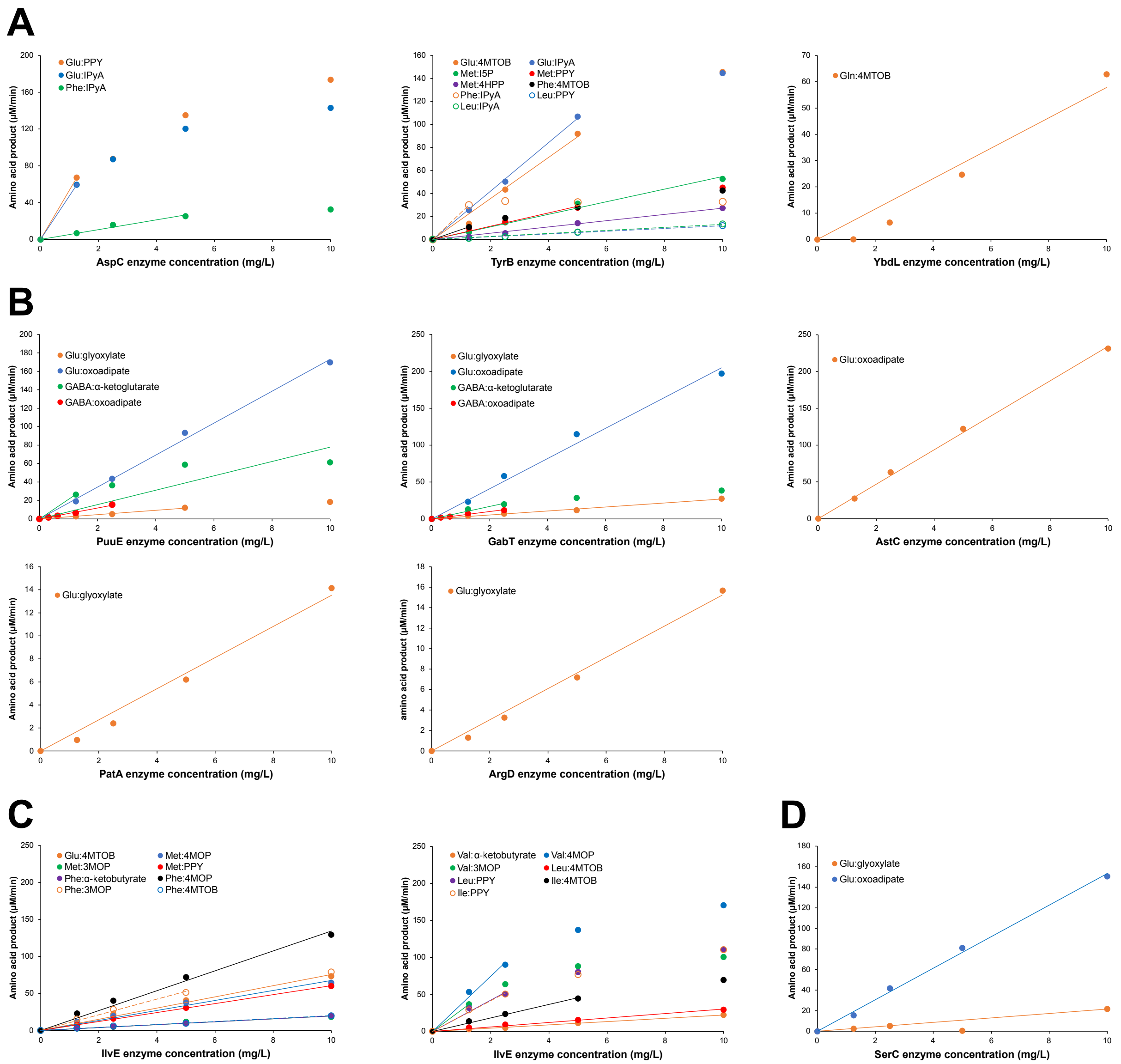


**Supplementary fig. S2. Validation of enzyme activity using HPLC-based assay**

AT activities with different combinations of amino donor:keto acceptor substrates detected from the LC-MS-based substrate mapping (**Fig. 2**) were validated using the HPLC-based enzyme assays. Amino acid products were detected by HPLC in the presence of indicated **(A)** Class I, **(B)** Class II, **(C)** Class III, or **(D)** Class IV EcAT enzymes at concentration of 0, 1.25, 2.5, 5, and 10 µg/mL, except for PuuE and GabT concentrations for the GABA:oxoadipate AT assay being 0, 0.3125, 0.625, 1.25, and 2.5 µg/mL.

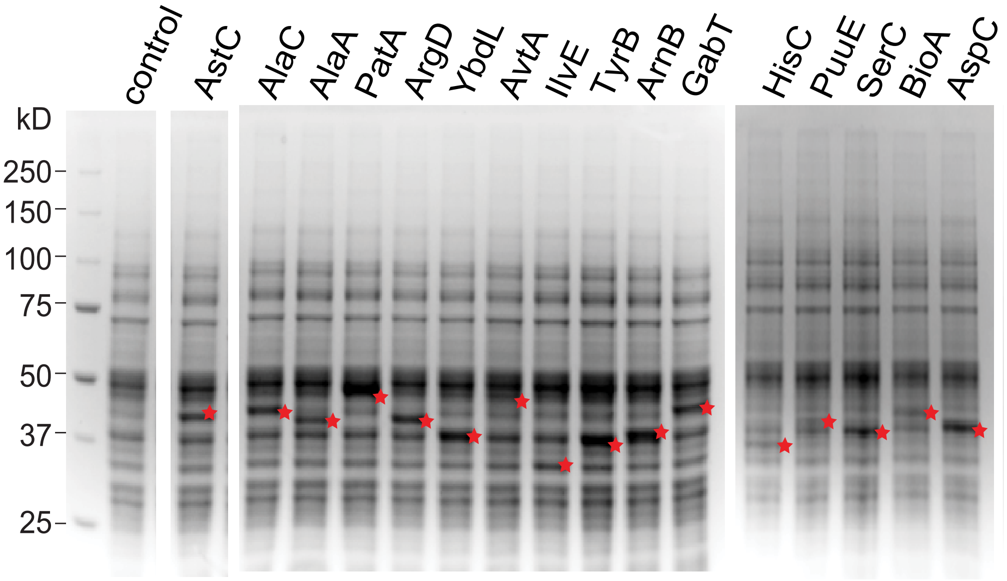


**Supplementary fig. S3. Sixteen *E. coli* enzymes synthesized by cell-free translation to be used as standards of the quantitative MS proteomics.**

The wheat cell-free protein synthesis produced 16 EcAT enzymes, which were used as quantitation standards for MS proteomics. The synthesized enzymes were detected together with wheat endogenous proteins by SDS-PAGE. Filled red stars indicate the synthesized enzymes.

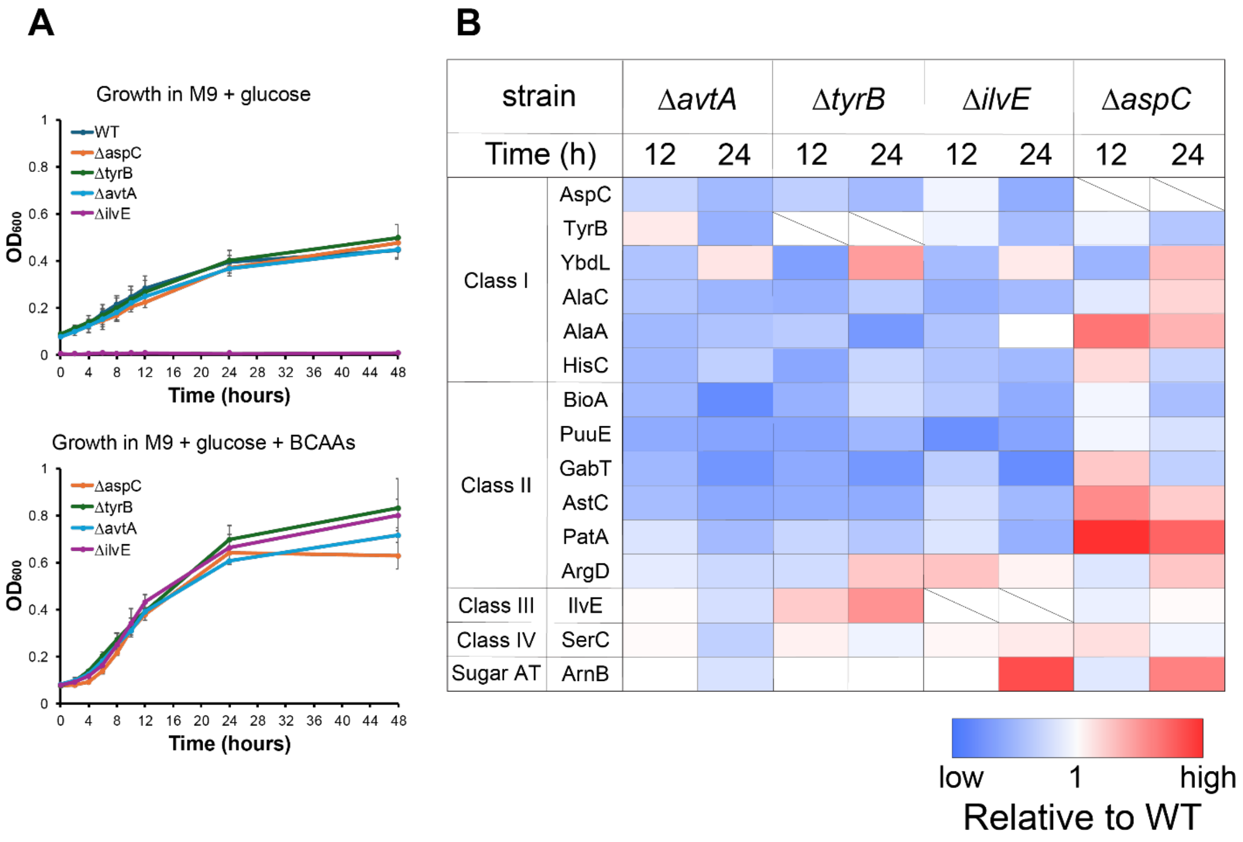


**Supplementary fig. S4. Intracellular changes in EcAT protein levels.**

**(A)** The growth curves of four single *EcAT* gene deletion strains in M9 medium with and without branched chain amino acids (BCAAs). The Δ*ilvE* mutant did not grow in the M9 media without BCAAs. No statistical difference was observed among genotypes, except for the lower $OD_{600}$ of Δ*aspC* at 48 hours in M9+BCAA media. Each data point is mean ± SD (n=3). **(B)** The relative abundance of EcATs in a single *EcAT* mutant to the wild-type from the representative dataset (n=1) is shown as the color code. The enzymes encoded by the deleted genes were marked with slashes.

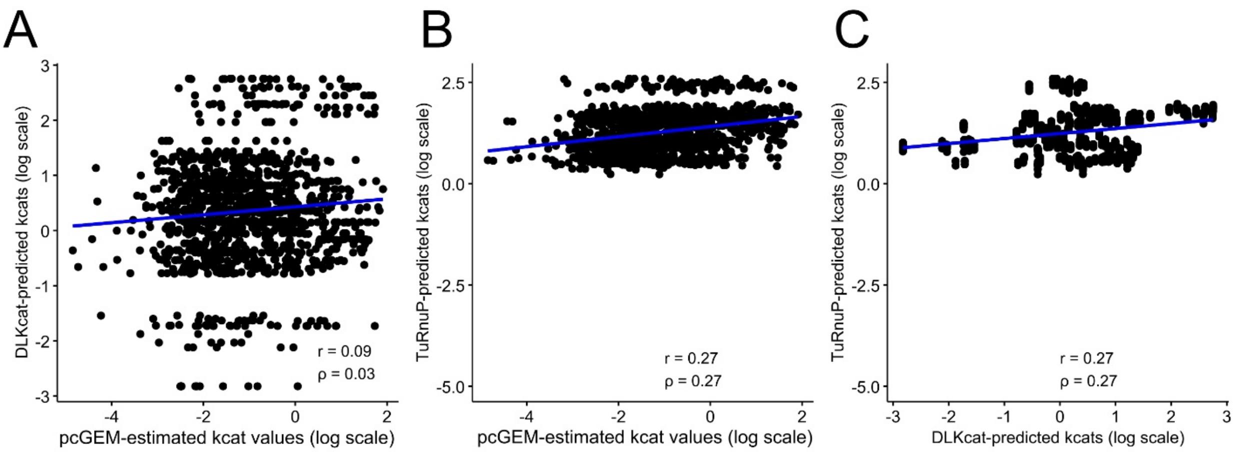


**Supplementary fig. S5. Comparison of estimated $k_{cat}$ values from the pcGEM to predictions of deep learning models**

The predicted $k_{cat}$ values of two deep learning models, DLKcat **(A)** and TurNuP **(B)**, presented low correlation to estimated $k_{cat}$ values using the protein-constrained eciML1515-EcAT model. The catalytic rates from the two deep learning models result in a low Pearson and Spearman correlations, of 0.09 and 0.03, respectively, for DLKcat; and 0.27 for both Pearson and Spearman correlations for TurNuP. TurNuP compared to DLKcat **(C)** yielded the same correlations of 0.27 for both Pearson and Spearman correlations. Pearson correlation is denoted by r and Spearman correlation is denoted by ρ, respectively.

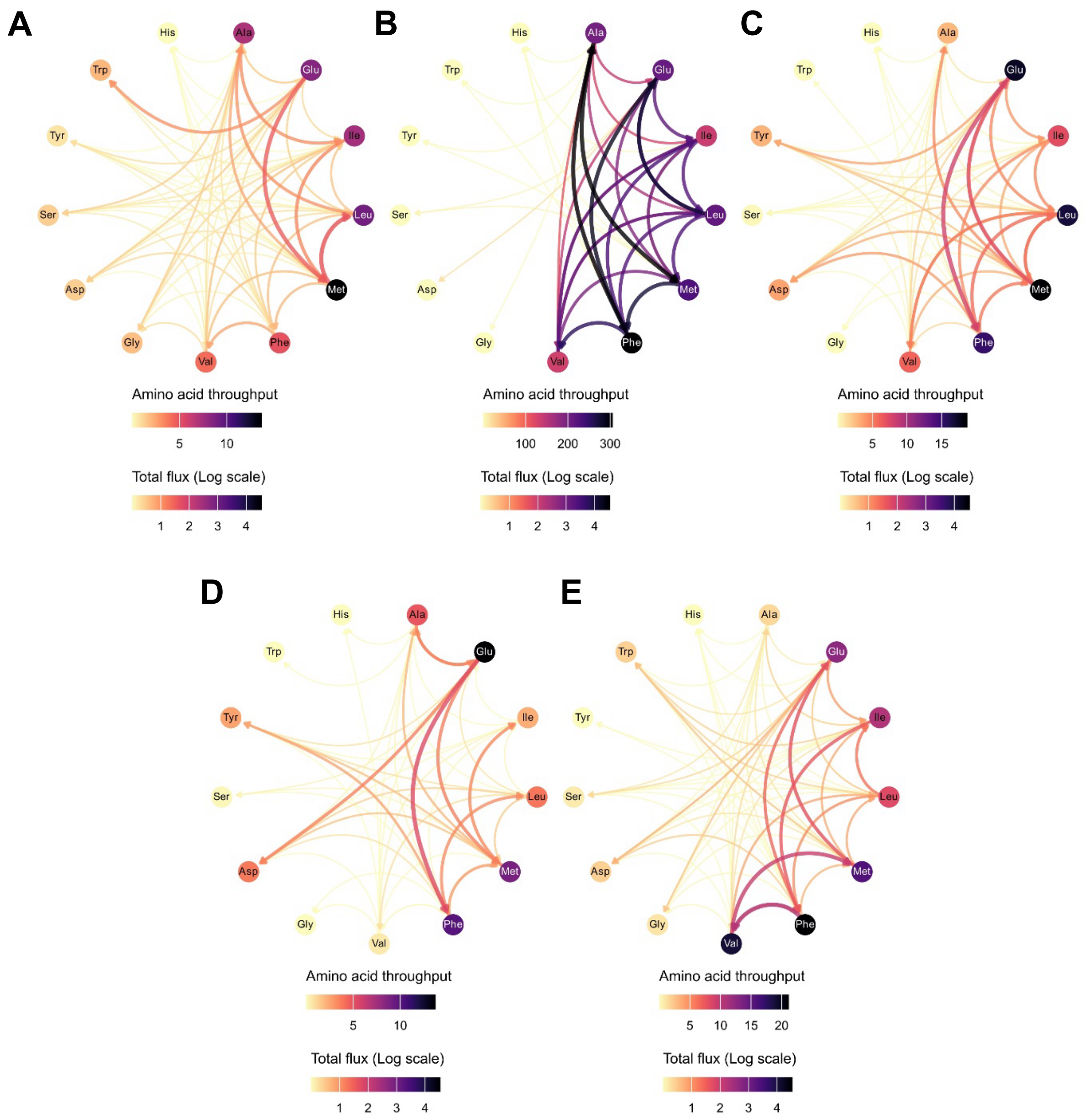


**Supplementary fig. S6. Mapping of nitrogen metabolism across strains.**

**(A)** Amino acid conversion graph for the WT strain. The total flux (represented in log scale) is the sum of metabolic fluxes through all reactions for a specific substrate conversion catalyzed by different enzymes across strains. Amino acid throughput represents the sum of the in-strength (sum of inward edge weights) and the out-strength (sum of outward edge weights) of all edges attached to that node. **(B)** Amino acid conversion graph for the Δ*aspC* strain. **(C)** Amino acid conversion graph for the Δ*tyrB* strain.**(D)** Amino acid conversion graph for the Δ*avtA* strain. **(E)** Amino acid conversion graph for the Δ*ilvE* strain.

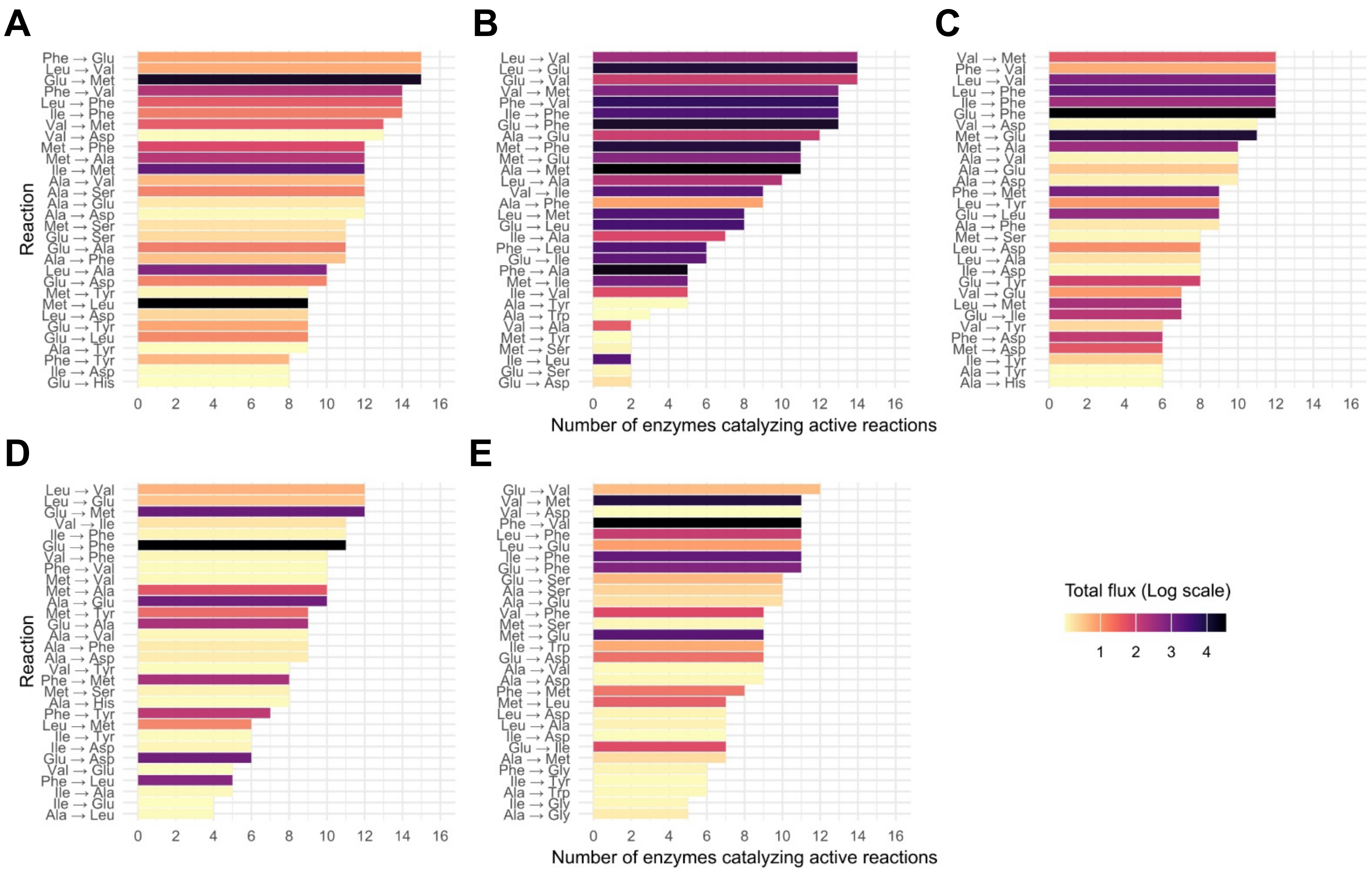


**Supplementary fig. S7. Enzyme activity counts across strains.**

**(A)** Enzyme activity counts in the WT strain. The total flux (represented in log scale) is the sum of metabolic fluxes through all reactions for a specific substrate conversion catalyzed by different enzymes across strains. For each substrate conversion reaction, multiple enzymes could be activated to catalyze the corresponding substrate conversion. **(B)** Enzyme activity counts in the Δ*aspC* strain. **(C)** Enzyme activity counts in the Δ*tyrB* strain. **(D)** Enzyme activity counts in the Δ*avtA* strain. **(E)** Enzyme activity counts in the Δ*ilvE* strain.

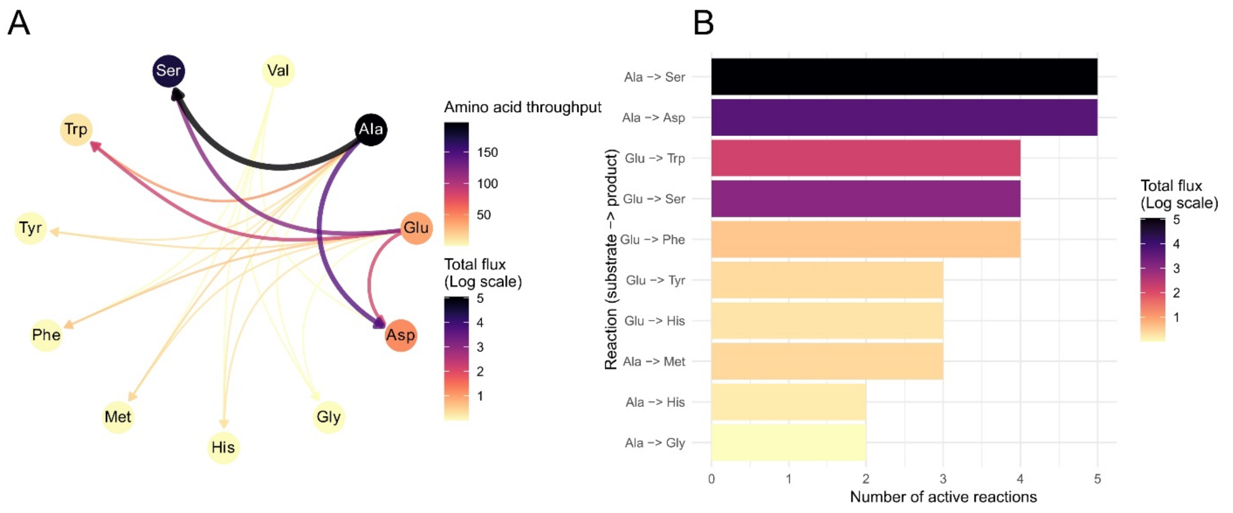


**Supplementary fig. S8. Mapping of nitrogen metabolism using the conventional GEM iML1515.**

For each substrate conversion reaction, multiple enzymes could be activated to catalyze that substrate conversion. **(A)** The reaction flux (represented in log scale) is the sum of metabolic fluxes through all reactions for a specific substrate conversion catalyzed by different enzymes across all strains. Amino acid throughput represents the total strength of the node, which is the sum of the in-strength (sum of inward edge weights) with the out-strength (sum of outward edge weights) of all edges attached to that node. **(B)** Active reactions are counted across all simulated conditions (WT and single-gene deletion strains). For iML1515, this is encoded solely in the GPR rules of each substrate conversion reaction. Since it does not consider enzyme constraints, the predictions rely on reaction stoichiometry.

**Supplementary Data:**

**Supplementary data S1. The list of *E. coli* AT enzyme genes and protein products.** This list contains information of the enzymes prepared in this study and the corresponding genes, including enzyme names, accession numbers, AT class, purity and concentration of expressed enzymes, amino acid and nucleotide sequences, and vector plasmid for gene synthesis.

**Supplementary data S2. % conversion of keto acid substrates to amino acid products as shown in Fig. 2A.** This dataset contains the mean values of two replicate experiments of 16 EcAT substrate mapping (Supplementary fig. S1) used to create the heatmap shown in Fig, 2A.

**Supplementary data S3. Primers used in this study**. This data lists primers used in this study, including primer name, target genes, nucleotide sequences, vector plasmid, and cloning method.

**Supplementary data S4. *E. coli* strains used in this study**. This data contains information of *E. coli* strains used in this study, including strain name, relevant genotypes, and source.

**Supplementary data S5. List of reactions and their associated $k_{cat}$values from the different parameterization approaches.** This dataset contains the AT reactions and the corresponding $k_{cat}$values predicted from multiple models. The AT reactions, the corresponding AT enzyme genes, and predicted $k_{cat}$ values using DLKcat, TurNuP, eciML1515-EcAT, and Retrained TurNuP.
